# Combinatorial CRISPR screening reveals functional buffering in autophagy

**DOI:** 10.1101/2020.07.28.201152

**Authors:** Valentina Diehl, Martin Wegner, Paolo Grumati, Koraljka Husnjak, Simone Schaubeck, Andrea Gubas, Varun Jayeshkumar Shah, Felix Langschied, Alkmini Kalousi, Ingo Ebersberger, Ivan Dikic, Manuel Kaulich

## Abstract

Functional genomics studies in model organisms and human cell lines provided important insights into gene functions and their context-dependent role in genetic circuits. However, our functional understanding of many of these genes and how they combinatorically regulate key biological processes, remains limited. To enable the SpCas9-dependent mapping of gene-gene interactions in human cells, we established 3Cs multiplexing for the generation of combinatorial gRNA libraries in a distribution-unbiased manner and demonstrate its robust performance. The optimal number for combinatorial hit calling was 16 gRNA pairs and the skew of a library’s distribution was identified as a critical parameter dictating experimental scale and data quality. Our approach enabled us to investigate 247,032 gRNA-pairs targeting 12,736 gene-interactions in human autophagy. We identified novel genes essential for autophagy and provide experimental evidence that gene-associated categories of phenotypic strengths exist in autophagy. Furthermore, circuits of autophagy gene interactions reveal redundant nodes driven by paralog genes. Our combinatorial 3Cs approach is broadly suitable to investigate unexpected gene-interaction phenotypes in unperturbed and diseased cell contexts.

## INTRODUCTION

Combinatorial gRNA expression (gRNA multiplexing) for related or orthogonal CRISPR applications enable the comprehensive characterization of genetic interactions in human cells. Several methods are available to support the generation of combinatorial gRNA expression systems, 1) restriction enzyme-based^1–4^, 2) golden-gate assembly^5–8^, 3) gateway-dependent^9,10^, as well as 4) recombination-dependent gRNA cloning techniques^11^. These systems are widely used to clone gRNA sequences in combination with RNA polymerase III promoters, resulting in arrayed gRNA-expression cassettes. In contrast, RNA polymerase II promoters generate RNA transcripts that can contain multiple gRNA sequences, although these transcripts require post-transcriptional processing to yield functional gRNA sequences and are currently limited to Cas12 applications^12,13^. The most widely used Cas-nuclease thus far is SpCas9, though a cloning-free gRNA multiplexing concept for Cas9 gRNAs is currently lacking, because repetitive and homologous sequences are unstable in lentiviral vectors^14–16^, rendering them less suited for large-scale combinatorial screening.

Concomitant mutations in two genes can yield unexpected phenotypes with respect to each gene’s individual phenotype^17^. Synthetic lethality as the most extreme combinatorial phenotype has clinical applications and is under therapeutic exploration^18–20^. A prominent example of a synthetic lethal gene pair with clinical relevance is poly (ADP-ribose) polymerase (PARP) inhibition in the context of defective *BRCA1* or *BRCA2* genes^15,21,22^. Additionally to this DNA-damage repair-related example, synthetic lethal interactions have been identified in combinatorial gRNA CRISPR screens, including the apoptotic genes *BCL2L1* and *MCL1* or *BCL2L1* and *BCL2L2*^16,23,24^, the mitogen-activated protein kinases 1 and 3 (*MAPK1* and *MAPK3*)^1^, as well as the PIP_3_ phosphatase *PTEN* and the mammalian target of rapamycin *MTOR*^25,26^. To perform pairwise hit calling in CRISPR screens, two methods are currently established: 1) a variational Bayes approach (GEMINI)^27^, and 2) the difference of expected to measured log_2_-fold-changes (dLFC) in which the expected log_2_-fold-change of a gRNA combination is the sum of the log_2_-fold-changes of each individual gRNA when partnered with control gRNAs^16,24^. These computational approaches have been applied to negative CRISPR screens. However, with combinatorial gRNA CRISPR screens being mostly performed in drop-out conditions, we lack knowledge of their performance for Cas9-based combinatorial gRNAs in positive or FACS-based phenotypic enrichment screens.

Bulk and selective autophagy are tightly regulated processes that target cellular material for lysosomal degradation and their misregulation culminates in abnormal cell growth and cell death with implications in various human diseases^28,29^. Rationally-engineered fluorescent reporter systems in combination with high-throughput CRISPR screens facilitated the systematic categorization of genes based on their essentiality in autophagy. As such, the transmembrane protein 41B, as well as the ubiquitin-activating enzyme UBA6, and the hybrid ubiquitin-conjugating enzyme/ubiquitin ligase BIRC6 were recently identified as important players in the autophagy network^30–35^. Despite the mapping of essential genes, CRISPR screens paired with fluorescent autophagic reporters revealed the stress-dependent regulation of autophagy-related genes^36–38^, and provided mechanistic insights into the gene specificity in bulk and selective autophagy by identifying PARKIN regulators and the ANT complex as essential components for mitophagy^39,40^. In combination with proximity biotinylation-coupled mass spectrometry, CRISPR screens also contributed to a spatial proteogenomic understanding of PARK2-dependent mitophagy^41^. Thus, unbiased CRISPR approaches coupled to fluorescent reporters and mass spectrometry are valuable approaches to identify vulnerabilities in bulk and selective autophagy for the treatment of neurodegenerative diseases and cancer^42–44^.

Here, we add to our previous work and describe the SpCas9-based 3Cs multiplexing technology for the generation of dual combinatorial (multiplexed) gRNA libraries with diversities of up to several hundreds of thousands of gRNA combinations^45^. In addition to identifying critical technical parameters, we generated a combinatorial library targeting the human autophagy network and performed a fluorescent reporter-based FACS enrichment screen to identify hitherto uncharacterized single genes and gene interactions essential for autophagy. Our 3Cs multiplexing technology is widely applicable and can serve as a general tool for the identification of context-dependent gene interactions at any scale.

## RESULTS

### 3Cs multiplexed Cas9 gRNA libraries

To expand on our 3Cs technology and enable Cas9 gRNA multiplexing, we generated a lentiviral Cas9-gRNA expression plasmid (pLenti-Multiplex) by placing a human 7SK (h7SK) promoter upstream of a previously engineered Cas9-tracrRNA followed by a human U6 (hU6) promoter upstream of a wildtype Cas9-tracrRNA sequence^46,47^. Both gRNA cassettes contain a gRNA placeholder sequence encoding for I-CeuI or I-SceI restriction enzyme sites, respectively (Figure 1A)^45^. Furthermore, in addition to a puromycin selection cassette, the plasmid contains an f1 bacteriophage origin of replication sequence in sense direction, supporting the CJ236 bacteria and M13KO7 bacteriophage-dependent generation of dU-containing single stranded (ss) DNA (Figure 1A, Supp. Figure 1A). In contrast to single gRNA 3Cs reactions, 3Cs multiplexing is performed in the presence of two gRNA-encoding oligonucleotide pools of which each contains unique 5’ and 3’ homology sequences for annealing to the h7SK or hU6 gRNA cassettes, thereby generating hetero-duplex dU-containing double stranded (ds) DNA that contains all possible gRNA combinations of the two oligo pools (Figure 1A). A coverage-based electroporation into *dut/ung*-positive bacteria results in template-strand depletion and combinatorial gRNA-containing dsDNA. To test this concept, we designed two gRNA-encoding oligo pools each containing 50 gRNA sequences targeting GFP (pool-1) or mCherry (pool-2) and used them to generate a combinatorial gRNA library, in addition to the respective single gRNA libraries (Figure 1B). Based on the typical three-band pattern of 3Cs dsDNA^45,48^, we observed similar yields and quality of dU-containing hetero-duplex dsDNA after applying two pools in a single 3Cs reaction when compared to either pool alone (Supp. Figure 1A). Digesting the final libraries with I-CeuI and I-SceI enzymes confirmed the exclusive presence of gRNA-containing plasmids (Figure 1C). Paired-end next-generation sequencing (NGS) revealed all three libraries to be complete with distribution skews of 1.56, 1.17, and 1.53 for GFP, mCherry, and GFP+mCherry, respectively (Figure 1D, Supp. Figure 1B-C). Most importantly, when packaged into lentiviral particles and transduced into GFP and mCherry-expressing hTERT-RPE1(Cas9) cells, the combinatorial GFP+mCherry library induced the simultaneous depletion of GFP and mCherry fluorescence while both single libraries selectively depleted either GFP or mCherry fluorescence (Figure 1E). Thus, 3Cs multiplexing generates tightly distributed combinatorial gRNA libraries that are functional in human cells.

**Figure 1.**
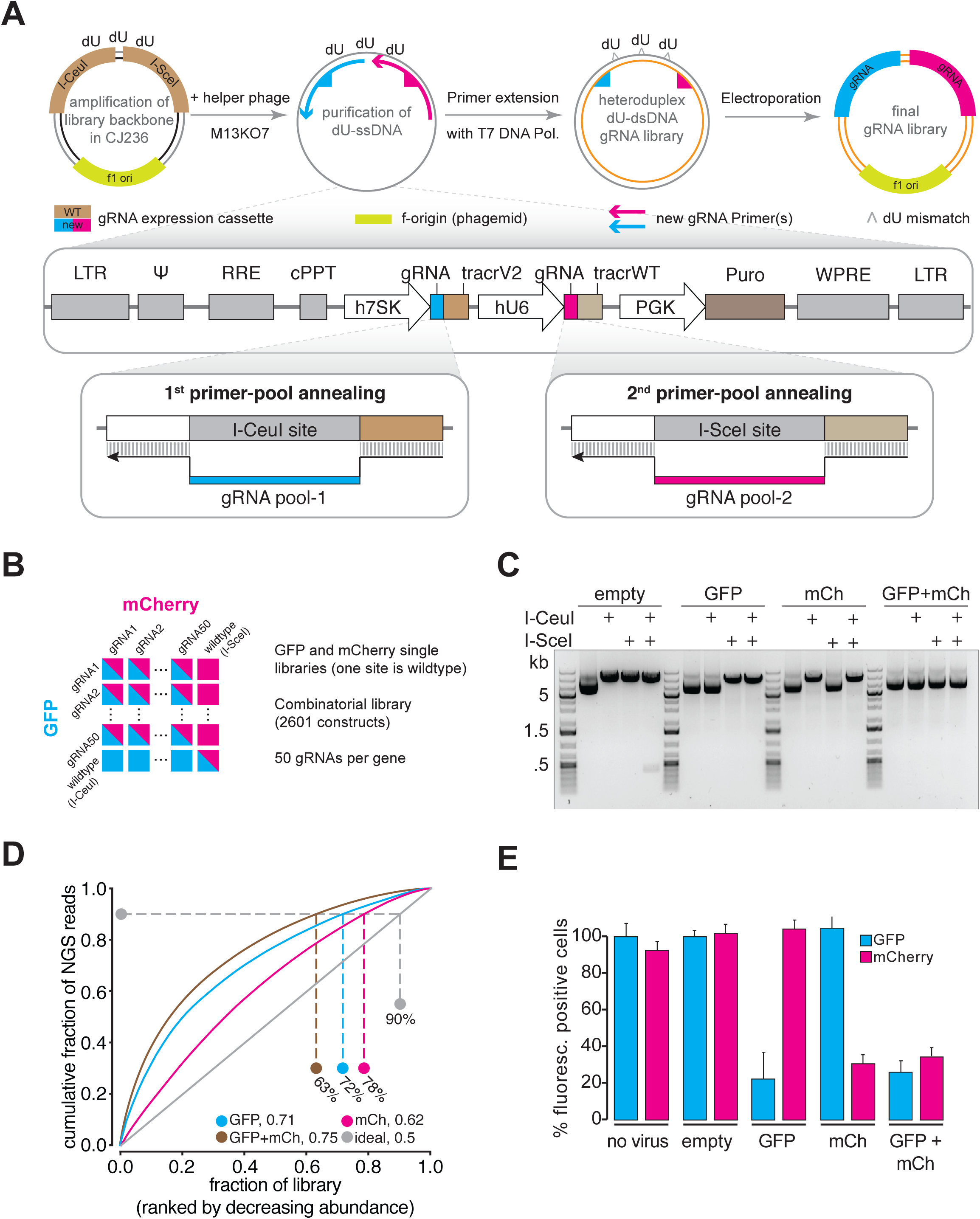
3Cs multiplexing for combinatorial CRISPR gRNA libraries. **A**) 3Cs multiplexing workflow. Two gRNA-encoding oligonucleotide pools are selectively annealed to one of the gRNA-expression cassettes of the dU-containing ssDNA template plasmid for T7 DNA Pol-dependent generation of heteroduplex dU-dsDNA. Amplification in dut/ung-positive bacteria amplifies the combinatorial gRNA plasmid. See Material and Methods for a more details. **B**) Cas9 GFP/mCherry multiplex library design. Combinatorial gRNA constructs target GFP and mCherry genes simultaneously; each gene is individually targeted by 50 gRNAs, both genes are simultaneously targeted with 2601 gRNA combinations. The single guide GFP-targeting library contains the wildtype gRNA placeholder in the hU6 cassette (mCherry), and vice versa. **C**) Gel-electrophoresis after analytical restriction enzyme digest of final 3Cs single and combinatorial libraries. **D**) Area-under-the-curve (AUC) determination of single and combinatorial library representation. As a reference, a perfectly distributed library (ideal) is shown in grey. Percentages indicate a library’s representation at 90% of cumulative reads. AUC values are indicated next to each library’s identifier. **E**) FACS analysis of GFP- and mCherry-positive hTERT-RPE1 cells after transduction with single or combinatorial libraries. Error bars represent standard deviations (SDs) over three biological replicates (n = 3).

### 3Cs multiplexing decouples sequence distribution and diversity

Current combinatorial gRNA libraries contain 9 to 18 gRNA pairs per gene-pair^3,15,24^, thus, a robust technology must generate large pairwise gRNA diversities without compromising reagent quality. We therefore investigated whether 3Cs multiplexing, similar to single gRNA 3Cs^45^, would decouple library quality from library size. To this end, we designed four oligonucleotide pools per gRNA cassette in which one, two, three, or four nucleotide positions within a non-human-target (NHT) gRNA sequence were randomized to mimic increasing combinatorial gRNA diversities (1N, 4*4=16 combinations; 2N, 16*16=256 combinations; 3N, 64*64=4,096, 4N, 256*256=65,536 combinations). NGS confirmed the libraries to be complete and evenly distributed with areas under the curve (AUC) between 0.59 and 0.69 (Figure 2A, Supp. Figure 2A-B). Notably, we identified the distribution skew to be very narrow with ranges from 1.1 (1N) to 1.49 (4N), values mostly unmatched even with single gRNA libraries (Supp. Figure 2A-B). Thus, 3Cs multiplexing is a highly robust method to generate combinatorial gRNA libraries with large sequence diversities.

**Figure 2.**
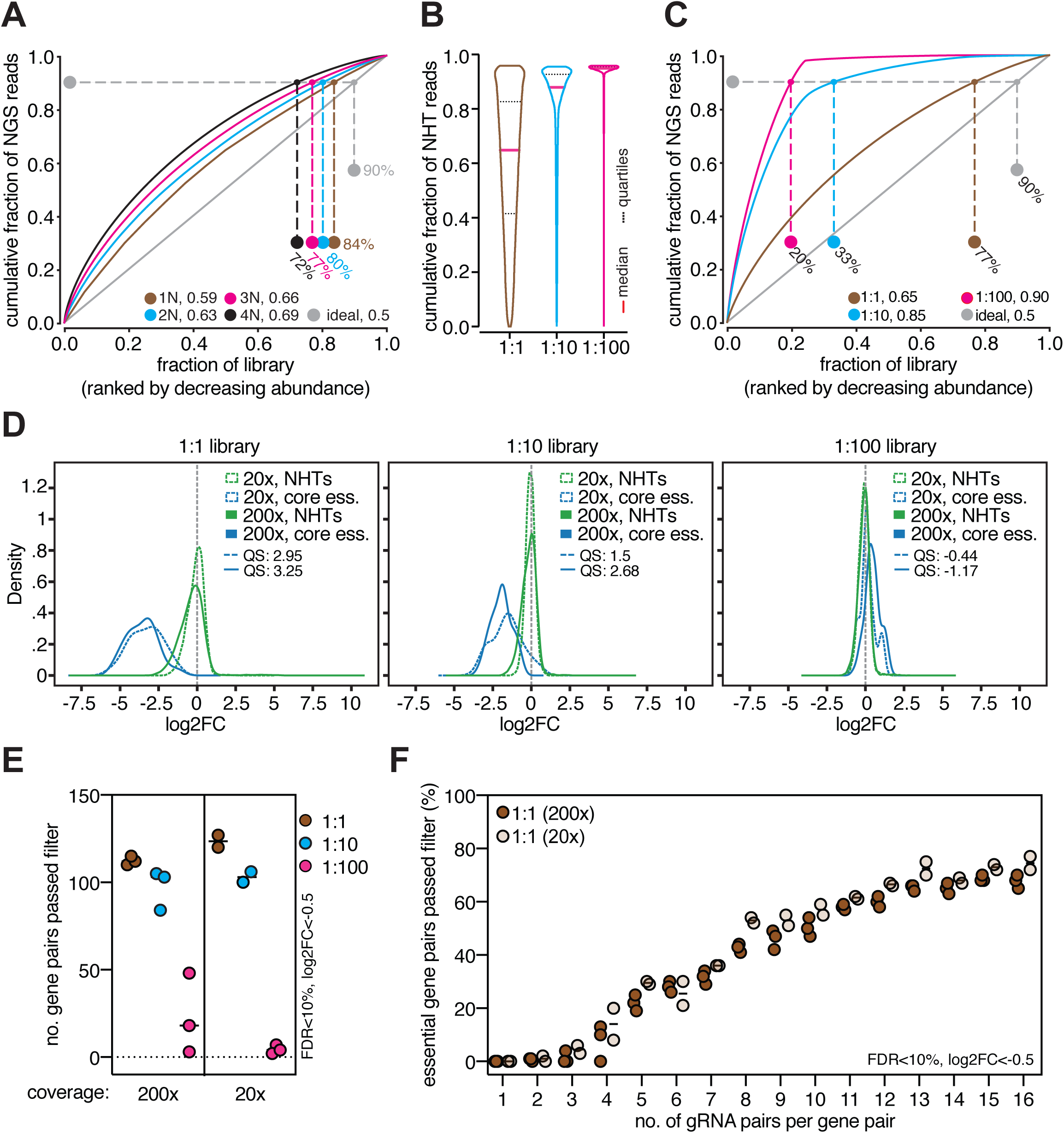
Library distribution skew dictates experimental scale and data quality. **A**) Area-under-the-curve (AUC) determination of the nucleotide-randomized libraries. As a reference, a perfectly distributed library (ideal) is shown in grey. Percentages indicate library representations at 90% of cumulative reads. AUC values are indicated next to each library’s identifier. **B**) Analysis of NHT-fraction in the biased 3Cs libraries. Median and quartiles of the distributions are shown as red straight and black dotted line, respectively. **C**) Area-under-the-curve (AUC) determination of the biased libraries. As a reference, a perfectly distributed library (ideal) is shown in grey. Percentages indicate a library’s representation at 90% of cumulative reads. AUC values are indicated next to each library’s identifier. **D**) Density plots showing the log2FC separation of combinatorial gRNA constructs targeting core essential genes (blue) and non-essential controls (green) in experimental coverages of 20x (dotted) and 200x (straight). QS: quality score. **E**) Analysis of the number of depleted gene-pairs detected with MAGeCK at the indicated FDR and log2FC cutoffs. **F**) Determination of the number of depleted gene-pairs at the indicated FDR and log2FC cutoffs from sub-sampled read-count tables of replicates for each coverage containing 1 to 16 randomly chosen gRNA pairs.

### Library distribution skew influences experimental scale

Higher replicate correlation was computationally demonstrated to correlate with smaller library distribution skew and higher library representation^49^. Thus, we aimed at experimentally investigating to what extent a combinatorial gRNA library’s distribution skew contributes to hit calling accuracy by generating combinatorial libraries with artificially distorted gRNA representations and screening them in different coverages. We selected a panel of 20 genes equally divided in 10 core essential (CE) and 10 tumor-suppressor genes (TSGs) and designed 4 gRNAs per gene. As internal controls, we choose 80 pre-validated NHT sequences^50,51^. We designed two 3Cs oligo pools: pool-1 contained CE and TSG-targeting gRNAs for h7SK and hU6 cassettes in equal ratios (2*80 gRNAs), while pool-2 consisted exclusively of NHT gRNA sequences in equal ratios for h7SK and hU6 cassettes (2*80 gRNAs). To generate artificially distorted library skews, we mixed both pools in equimolar ratios (1:1), and in ratios of increasing NHT sequence-molarity (1:10 and 1:100) and applied them to 3Cs multiplexing. NGS confirmed an increased fraction of NHT reads in the 1:10 and 1:100 libraries (Figure 2B, Supp. Figure 2C), and revealed an AUC value for the 1:1 library of 0.65, while the 1:10 and 1:100 libraries contained increased AUC values of 0.85 and 0.9, respectively (Figure 2C). Most importantly, library distribution skews increased from 1.2 (1:1) to 2.4 (1:10) and 13.46 (1:100) (Supp. Figure 2D), demonstrating that final 3Cs library quality is coupled to an oligo pool’s distribution.

Using these libraries in representations of 20- and 200-fold, we evaluated a total of 153,600 pairwise gRNA-knockouts in human hTERT-RPE1(Cas9) cells in biological replicates. Pairwise guide and gene-level counts correlated well for individual libraries and biological replicates (gene level, 20x - 1:1, Pearson r = 0.96; 1:10, Pearson r = 1; 1:100, Pearson r = 0.79-1; 200x - 1:1, Pearson r = 0.91-0.98; 1:10, Pearson r = 0.96-0.97; 1:100, Pearson r = 0.98-1; Supp. Figure 3A-B). We next analyzed the difference of log_2_ fold changes (log2FC) in our screens by means of separating essential genes from non-targeting controls and computed Cohen’s D statistics (quality score (QS)), a value recently introduced to quantify screen quality^52^. Quality scores for the 1:1 library in both tested coverages were very high with 2.95 and 3.25 for 20- and 200-fold, respectively (Figure 2D). However, a decline in screen quality appeared when library distribution skews increased above 2 (1:10 library, 20x - QS: 1.5, 200x - QS: 2.68; 1:100 library, 20x - -0.44, 200x - -1.17; Figure 2D).

We next assessed if higher experimental coverage was able to rescue wider library distribution skews by identifying essential gene pairs with MAGeCK analyses and cut-off filters set to FDR≤10% and log2FC<-0.5. Strikingly, the number of retrieved gene pairs per library distribution skew was nearly identical between 20- and 200-fold coverages (Figure 2E), suggesting that CRISPR libraries with narrow distribution skews can be screened at minimal experimental coverage without compromising data quality. Furthermore, these analyses also suggest that higher experimental coverage is likely insufficient to rescue wider library distribution skews. Therefore, a library’s distribution skew is a critical parameter that should be considered when designing combinatorial CRISPR libraries and screens.

### The optimal number of pairwise gRNAs for statistical gene-interaction calling

The number of gRNAs per gene has a profound impact on statistical hit calling^53,54^, thus we analyzed the concordance of essential pairwise genes identified across the two experimental coverages with increasing numbers of pairwise gRNAs. We down-sampled the 1:1 data sets and generated a series of read count tables containing between 1 and 16 randomly chosen gRNA pairs for each gene combination and performed MAGeCK analyses with cut-off filters at FDR≤10% and log2FC≤-0.5^53,54^. A total of 100 essential gene interactions exist within both libraries (10*10 essential genes), and as expected, 16 pairwise gRNAs per gene interaction consistently retrieved the largest number of statistically significant essential gene interactions (200x - 70%, 20x - 72%) (Figure 2F). However, performance differences became evident in the down-sampled libraries containing up to 4 pairwise gRNAs with 13% and 20% paired gene interactions identified, for 200 and 20-fold library representations, respectively (Figure 2F). Strikingly, the performance increased noticeably in both coverages up until 13 pairwise gRNAs from which on the performance plateaued (Figure 2F). These results are consistent with previous observations for single gRNA SpCas9-dependent CRISPR screens, in which 4 to 6 gRNAs per gene have been identified for robust statistical hit calling^55^. Assuming the generation of gRNA number-balanced libraries, our observations identify 16 pairwise gRNAs (4*4) as the optimal number for statistical gene-interaction calling.

### Single and inherent-single gRNA enrichment screens are complementary

Pairwise gRNA screens have been applied to phenotypes related to cell viability (drop-outs), but their performance in enrichment-based screens remains elusive. To address this question, we identified parameters to investigate pairwise gene interactions in the self-degrading process of human autophagy by LC3-coupled fluorescent-dependent enrichment screening. We used the established GFP-LC3-RFP reporter construct to generate a monoclonal hTERT-RPE1 reporter cell line and confirmed reporter functionality by mTOR inhibition (Torin1 treatment) alone and in combination with the autophagy inhibitor Bafilomycin (Supp. Figure 4A)^56^. Next, we assembled a literature-curated list of 192 “extended autophagy” genes that included, among others, autophagy-related genes (ATGs), autophagy receptors, as well as transcription factors and deubiquitinases, plus 7 core essential genes as negative controls. To generate a combinatorial gRNA library, we designed a second oligo pool, named “core autophagy”, consisting of 64 commonly considered core autophagy genes (Figure 3A, core autophagy, Supp. Table 1). For each gene, we designed 4 gRNA sequences using rule set 2 and added 10% NHT sequences to each pool^51^, resulting in a total of 876 gRNAs in the “extended autophagy” pool and 282 gRNAs in the “core autophagy” pool, together generating 247,032 pairwise gRNAs (Figure 3A). We applied the extended autophagy pool alone and in combination with the core autophagy pool to 3Cs reactions and generated the single gRNA and pairwise gRNA libraries, respectively. NGS revealed narrow distribution skews (single library, skew ratio=1.22; multiplex library, skew ratio=1.69) and AUC values of 0.65 and 0.68 (Supp. Figure 4B-D), supporting our previous observation that 3Cs multiplexing is a highly robust method for the generation of combinatorial gRNA libraries.

**Figure 3.**
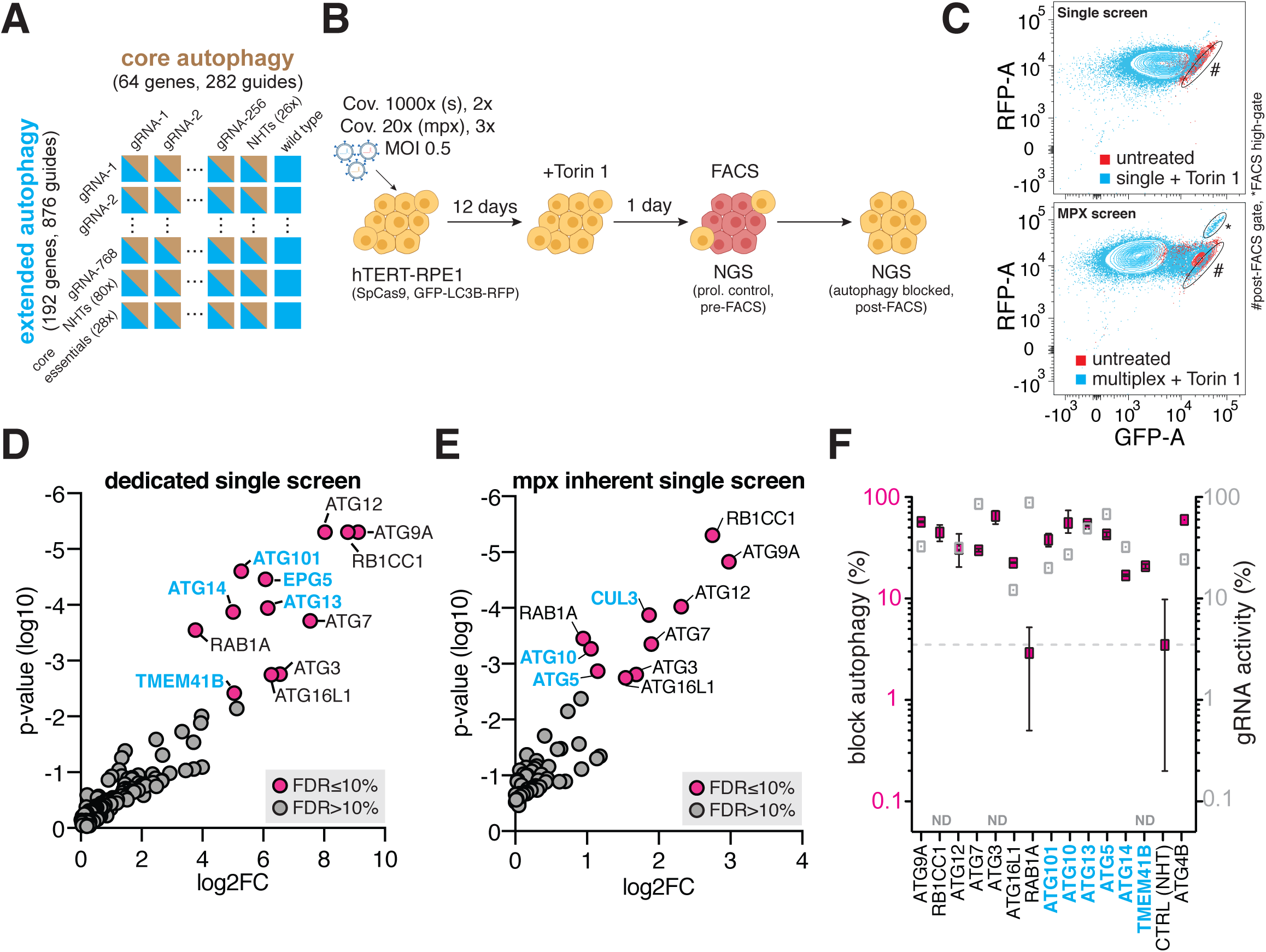
Single and multiplex-inherent single gRNA enrichment screens are complementary. **A**) Cas9 autophagy single and multiplex library design. Combinatorial gRNA constructs target extended and core autophagy genes; each gene is targeted by 4 gRNAs, 192 extended autophagy genes and 64 core autophagy genes with 10% of NHT controls per cassette generating 247,032 gRNA combinations. Single gRNA extended autophagy library contains the wildtype gRNA placeholder in the hU6 cassette (core autophagy). **B**) Single (s) and combinatorial (mpx) autophagy screening workflow. Cov.: coverage (library representation); MOI: multiplicity of infection; FACS: fluorescence-activated cell sorting; NGS: next-generation sequencing. **C**) Cell sorting representations of single and combinatorial (MPX) screen of post-FACS gate (#) and high-gate (*). **D**) MAGeCK analysis of dedicated single gRNA autophagy screen of pre- and post-FACS samples with hit genes in red when matching cutoff criteria of FDR≤10% and p-value≤0.05. Screen-selective hits in blue. **E**) MAGeCK analysis of multiplex-inherent single gRNA autophagy screen of pre- and post-FACS samples with hit genes in red when matching cutoff criteria of FDR≤10% and p-value≤0.05. Screen-selective hits in blue. **F**) Analysis of single hit genes derived from D) and E) in arrayed autophagy blockage validations (red). Evaluation of gRNA activity by TIDE analysis (grey). Error bars represent standard error of mean (SEM) over three biological replicates per autophagy blockage (n=3). ND: not determined.

To benchmark our screening approach, we first applied the single autophagy library to screen for genes essential for Torin1-induced autophagy (Figure 3B). In order to reduce the number of false-positive hits, we chose stringent FACS-gating criteria that only enriched cells in which autophagy was completely blocked (Figure 3C, single screen). Importantly, proliferative effects as a consequence of gene knockouts are a source of false-positive hit calling in enrichment screens, especially in TP53-positive cells. Thus, we collected a coverage-based cell sample at the day of FACS sorting to correct for positive or negative proliferative effects (Figure 3B). Guide RNA and gene level correlations between pre- and post-FACS samples were high among biological replicates (guide level, Pearson r=0.87; gene level, Pearson r=0.97; Supp. Figure 5A-B), thus we applied MAGeCK and retrieved 12 significantly enriched genes with log2FC between 3.76 and 9.12 with FDR≤10% (Supp. Table 2). Importantly, our benchmark screen successfully retrieved previously identified core autophagy genes required for mTOR-induced autophagy and correctly assigned their function to bulk autophagy by means of a positive log2FC between pre- and post-FACS conditions (Figure 3D)^32,33,35,39,40,57,58^.

Next, we dramatically scaled up to perform the autophagy multiplex screen at a 20-fold coverage at pre- and post-FACS time points, investigating a total of 1,235,160 pairwise gRNAs. Guide RNA and gene level correlations between pre- and post-FACS samples were high among biological triplicates: gRNA level, pre-FACS, Pearson r=0.99; post-FACS, Pearson r=0.58-0.69; gene level, pre-FACS, Pearson r=0.99; post-FACS, Pearson r=0.77-0.84 (Supp. Figure 5A, C-D). During the course of FACS sorting, we noticed a population of cells that was absent from single gRNA transduced cells and extended our sorting strategy to also investigate cells from this “high-gate” (Figure 3C). We first assessed the nature of the high-gate cell population and identified the majority of reads to account for ATG4B-dependent gRNA pairs (post-FACS 1, 86.94%; post-FACS 2, 78.03%; post-FACS 3, 75.53%) (Supp. Figure 5E). To permit its function, ATG4 cleaves the LC3-reporter and paired ATG4 gene knockouts likely interfere with reporter functionality and served as additional controls to validate our library and screening strategy^56^.

Within the combinatorial gRNA library, four sets of gRNA interactions occur, 1) control with control (NHT:NHT), 2) extended autophagy with control (gene:NHT), 3) control with core autophagy (NHT:gene), and 4) extended autophagy with core autophagy (gene:gene) (Figure 3A). Thus, we next assessed the concordance of identified single genes essential for autophagy induction between our single autophagy benchmark screen and the extended autophagy gRNAs paired with control guides in the combinatorial screen. We applied MAGeCK to compare pre- and post-FACS time points and identified 10 significantly enriched genes with log2FC between 1.53 and 2.75 and FDR≤10% (Figure 3E). Screen concordance was high with 7 jointly identified genes, though each experiment successfully identified unique genes essential for bulk autophagy induction (Figure 3D-E, unique hits in blue-bold). Importantly, the jointly identified genes ATG9A, RB1CC1, ATG12, ATG7, ATG3, and ATG16L1 as well as the unique hit genes ATG101, ATG10, ATG13, ATG5, ATG14 and TMEM41B could be successfully validated by targeting each gene with a newly designed single gRNA, for which gRNA performance during the course of validation was quantified by TIDE analysis (Figure 3F). Screen concordance could be further improved to 11 jointly identified genes by filtering for p-values below 0.05 and simultaneously relaxing the FDR, although the number of uniquely identified genes per screen increased accordingly to 8 and for dedicated single and inherent single screens, respectively (Supp. Figure 6A-B). This demonstrates that dedicated single and inherent-single autophagy screens are complementary and that either one alone was insufficient to identify all true-positive hits.

### Synergistic gene pairs essential for autophagy

Targeting single genes with two gRNAs has been shown to improve knockout rates^59^, we therefore investigated the consistency of this phenotype within our combinatorial data set. As expected, we repeatedly observed a higher log2FC for combinatorial-gRNA targeted genes (Supp. Figure 7A), particularly for genes that we identified and validated as essential for autophagy (Supp. Figure 7B). Next, we assessed pairwise gene interactions essential for autophagy induction by applying MAGeCK to pre- and post-FACS samples. This identified a total of 3645 significant gene pairs, of which 187 gene pairs accounted for pairs in which both genes were jointly or uniquely identified as single essential for autophagy (Figure 4A, D). 3245 gene pairs accounted for gene pairs in which one or the other gene was identified as being essential for autophagy (Figure 4B, D), and 213 gene pairs consisted of genes for which neither partner gene was identified as being essential for autophagy (Figure 4C, D). Interactions between the majority of identified gene pairs are driven by a single gene that is essential for autophagy and interactions between non-essential autophagy genes are rare, which is well in agreement with previous work that mapped genetic interactions in *Saccharomyces cerevisiae*^60^. To corroborate our findings, we set up arrayed dual gene-targeting validations in which either one gene or both genes were essential for autophagy induction. As expected, targeting two essential autophagy genes consistently increased the fraction of cells in which autophagy was blocked (Figure 4E), and pairing an essential autophagy gene with an autophagy-related but non-essential gene also increased the fraction of cells in which autophagy was blocked (Figure 4F). Interestingly, among the non-essential:non-essential hits, we identified ULK1, AMBRA1, WIPI2, and BECN1 in combinations with control gRNAs, suggesting these genes to be essential for autophagy on their own, a conclusion supported by the literature^33,57^. They have likely missed our attention in the single gRNA CRISPR screens due to their relatively mild phenotype compared to the other autophagy essential genes. Indeed, in arrayed single gRNA validations, the depletion of ULK1 and WIPI2 alone blocked autophagy, although the phenotype was milder when compared to ATG9A depletion that we identified as a strong hit (Figure 4G). This suggests, in the context of hTERT-RPE1 cells and Torin1-induced autophagy, that gene-associated categories of phenotypic strengths exist in autophagy. Most interestingly, among others, we identified several gene pairs capable to block autophagy induction (Figure 4C). We noted several gene pairs in which one gene is either of the Ras-related protein family (RABs) as well as several pairs that included ATG2A. Among these interactions we identified ATG2A-ATG2B for which the validation in arrayed dual-gene knockout conditions confirmed the blockage of autophagy only when both genes are interfered with, experimentally verifying these genes as functional redundant homologues (Figure 4G).

**Figure 4.**
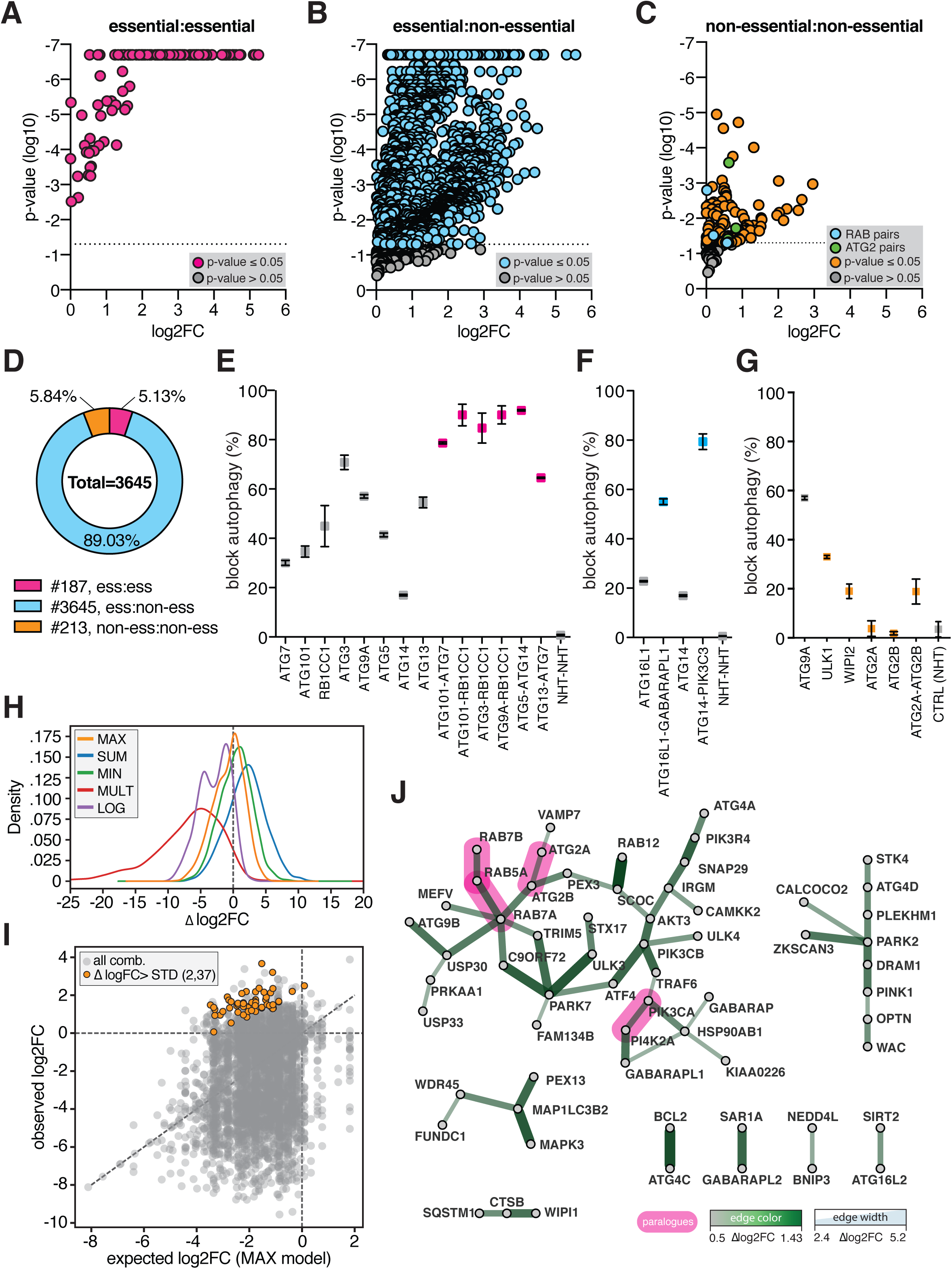
Paralogs are redundant in autophagy. **A**) MAGeCK analysis of combinatorial gRNAs targeting gene pairs in which both genes were identified as essential for autophagy of pre- and post-FACS samples. Hit gene pairs are shown in red when matching cutoff criteria of p-value≤0.05. **B**) MAGeCK analysis of combinatorial gRNAs targeting gene pairs in which one gene was identified as essential for autophagy of pre- and post-FACS samples. Hit gene pairs are shown in blue when matching cutoff criteria of p-value≤0.05. **C**) MAGeCK analysis of combinatorial gRNAs targeting gene pairs with neither gene being identified as essential for autophagy of pre- and post-FACS samples. Hit gene pairs are shown in yellow when matching cutoff criteria of p-value≤0.05. Ras-related protein family (RAB) and ATG2 gene pairs are shown in blue and green, respectively. **D**) Global view on identified gene pairs per category in percent. Ess: essential; non-ess: non-essential. Color code adapted from A) to C). **E-G**) Arrayed analysis of hit gene pairs and the induced blockage of autophagy per gene knockout of each category, color code of A) to C). Control genes and sequences are shown in grey. Error bars represent standard error of mean (SEM) over three biological replicates (n=3). **H**) Density plots of delta log2FC (Δlog2FC) analyses computed by MAX, SUM, MIN, MULT, and LOG models. **I**) Correlation between observed and expected log2FC values, derived from MAX model, for combinatorial gene-targeting. Data points above standard deviation and with p-values ≤0.05, derived from C), are highlighted in yellow, representing synergistic gene interactions in autophagy. **J**) Network analysis of synergistic autophagy gene pairs, derived form I). Edge color and width set to Δlog2FC values derived from MAX model. Edges and nodes of paralog gene pairs are highlighted in pink.

Four mathematical definitions of genetic interactions have been proposed previously (Product (MULT), Additive (SUM), Log (LOG), and Min (MIN))^17^. We also included a Max (MAX) definition and applied all models to identify combinatorial phenotypes that are surprising with respect to each gene’s single phenotype. Genetic interactions are rare^60^, we thus computed the deviation of observed paired-knockout phenotypes from their expectation (delta log2FC, Δlog2FC) and identified the MAX model to fit our combinatorial autophagy data set best (Figure 4H), especially when only considering genes that we identified to be essential for autophagy for which the deviation from the expectation is low (Supp. Figure 7C). In total, we identified 3665 genetic interactions that we filtered by significance (p≤0.05) and Δlog2FCs above their single standard deviation, resulting in a total of 57 (1.6%) high confidence synergistic gene pairs (Figure 4I, yellow dots, Supp. Table 3). To better depict relations among these gene interactions, we performed a network analysis and identified 5 network components (1, 36 nodes; 2, 10 nodes; 3, 5 nodes; 4, 3 nodes; and 5; 4 x 2 nodes) (Figure 4J). Most importantly, we identified functional redundancy in autophagosome assembly (ATG2A/B), phosphatidylinositol phosphorylation (PIK3CA/PI4K2A), phagosome-lysosome fusion (RAB7A/5A and RAB7B/5A), and selective autophagy (PARK2 with PLEKHM1 and CALCOCO2, OPTN with PINK1, and PARK7 with FAM134B) (Figure 4J). Furthermore, we identified DNA Damage-Regulated Autophagy Modulator Protein 1 (DRAM1)-dependent interactions with mitochondrial PINK1 and PARK2 for which the DRAM1-dependent induction of autophagy was shown to be dependent on mitochondrial protein synthesis inhibition^61^. Thus, our identified gene interactions likely resemble functional nodes within the human autophagy circuit in which redundancy exists and point towards gene paralogs as a critical factor in generating redundancy in autophagy.

## DISCUSSION

Several technologies are available to generate combinatorial gRNA-containing plasmids and libraries. They mostly depend on open plasmid DNA and PCR amplified gRNA-encoding oligonucleotide pools, resulting in cloning-artefacts and sequence biases. Our newly developed 3Cs multiplexing technology functions with single stranded template plasmids and oligonucleotide pools, thereby circumventing additional and unintended sequence representation dispersions.

Passaging coverage and CRISPR library distribution skew have recently been computationally predicted to be critical factors for data quality^49^. Indeed, our experimental analysis of combinatorial libraries with varying distribution skews, applied with different passaging coverages, supports this prediction and identifies a distribution skew of below 2 as a threshold enabling passaging coverages below 100-fold. Furthermore, we provide experimental evidence that combinatorial libraries with distribution skews above 2 should be screened with passaging coverages ≥200 to compensate for the library’s sequence dispersion. However, rescuing libraries with large distribution skews (≥10) by applying high passaging coverage is likely going to fail, as the sequence dispersion is too big to be compensated for due to sequence underrepresentation at screen initiation.

Combinatorial gRNA screens are more frequently performed to identify gene pairs essential for cell viability. However, as of today, no combinatorial gRNA screen in combination with pathway-specific fluorescence reporters has been performed. While the inherent single gRNA phenotypes appear to be sufficient in the context of cell viability screenings, we show that dedicated and inherent single gRNA screens are complementary for combinatorial enrichment screens. This is likely due to the strong enrichment of combinatorial gene phenotypes and the associated underrepresentation of single-gene effects. However, this can be compensated for by including a dedicated single gene screen when planning combinatorial enrichment screens. Furthermore, we note that genes associated with strong phenotypes are sufficiently identified by this approach, but genes causing milder phenotypes will likely miss attention. This obstacle, however, is circumvented by carefully examining gene-interactions in the class of phenotype-associated non-essential gene pairs. Indeed, we demonstrate that AMBRA1, ULK1, WIPI2, and BECN1 are essential for autophagy, even though both single gRNA screens failed to identify them.

Genome-wide or combinatorial CRISPR screens demand large numbers of cells. Thus, current efforts aim at minimizing cell culture demands by providing combinatorial-gRNA minimized CRISPR libraries^59,62^. Supporting this notion, we identify log2FC of dual-gRNA targeted genes to be larger than their single-gRNA targeted counterpart. Interestingly, this observation was limited to genes that we identified to be essential for autophagy, supporting the notion that dual-gRNA gene targeting also induces stronger phenotypes in enrichment screens and that minimized libraries will also be beneficial for these applications.

Functional buffering by paralogs has recently been shown to be largely absent from single gRNA CRISPR screens^63^, suggesting paralogs to contribute to network redundancy. Indeed, we identify paralogs as functional buffers in autophagy acting in autophagosome assembly (ATG2A/B), phosphatidylinositol phosphorylation (PIK3CA/PI4K2A), and phagosome-lysosome fusion (RAB7A/5A and RAB7B/5A). This is profoundly important when therapeutically targeting autophagy induction in cells that are dependent on high basal autophagy levels such as acute myeloid leukemia (AML) cells^64^. Lastly, we note that our analysis failed to identify functional buffering within the mammalian ATG8 family of proteins (LC3s and GABARAPs), as well as within the protein class of autophagy receptors (FUNDC1, SQSTM1, OPTN, PLEKHM1, PEX13, CALCOCO2, FAM134B). Furthermore, the lack of buffering gene interactions between ATG8s and autophagy receptor proteins, together, supports the notion of their cargo selectivity that prevents these genes to contribute to autophagy redundancy^65^.

## MATERIAL AND METHODS

### 3Cs multiplex template plasmid DNA and cloning

pLentiCRISPRv2 (Addgene: 98290) was enzymatically digested with AleI and BsiWI and gel purified to remove the hU6 gRNA- and SpCas9-expressing cassettes. Likewise, the combinatorial gRNA-expressing cassette of pKLV2.2 (Addgene: 72666) was digested with AleI and BsiWI. The 2030 bp fragment that encodes the combinatorial gRNA-expressing cassettes and a PGK promoter was gel purified and cloned into the reduced, purified backbone of pLentiCRISPRv2. In order to generate unique annealing homology for the 3Cs oligonucleotides and enable template plasmid removal, the h7SK promoter-associated tracrRNA was replaced by a previously engineered tracrRNA sequence (tracrRNA v2) and h7SK and hU6 promoter-associated gRNA cloning sites were modified to contain placeholder sequences encoding for I-CeuI and I-SceI homing endonuclease restriction sites, respectively^66^.

### 3Cs oligonucleotide design rules

All oligonucleotides that were used for multiplexed 3Cs gRNA library generation are listed in ‘DNA oligonucleotides’. DNA oligonucleotides were purchased from Sigma-Aldrich, from Integrated DNA Technologies (IDT) as single or pooled oligonucleotides in o-pool formats, and from Twist Bioscience as oligonucleotide pools.

To discriminate between h7SK and hU6 and enable exclusive annealing to only one expression cassette, the 3Cs oligonucleotides were designed with two specific homology regions flanking the intended 20-nt gRNA sequence for either the h7SK or hU6 expression cassettes. The 3Cs h7SK-oligonucleotides were 57 nts in length (Tm above 50°C) and matched the 3′ end of the h7SK promoter region and the 5′ start of the tracrRNA v2, while the 3Cs hU6-oligonucleotides were 59 nts in length (Tm above 50°C) and matched the 3′ end of the hU6 promoter region and the 5′ start of the tracrRNA v1 in the template plasmids.

### Generation of sequence distorted 3Cs libraries

For the generation of biased multiplex 3Cs gRNA libraries, two 3Cs oligonucleotide pools were designed for each expression cassette of the 3Cs multiplex template plasmid following the 3Cs oligonucleotide design rules. The first pool was composed of tumor suppressor and essential gene-targeting gRNAs (target pool), while the second pool only included non-human targeting gRNAs (control pool). To generate three different libraries that represent libraries of different quality regarding their distribution, the two oligonucleotide pools were mixed in a different ratio for each of the three libraries. For the first library, the target and control pool were mixed in a 1:1 ratio, to resemble an evenly distributed gRNA library. For the second library a 1:10 ratio of target to control pool was applied. The third library was generated with a 1:100 ratio, to resemble a library with highly underrepresented gRNA sequences. The mixed oligonucleotide pools were phosphorylated, annealed to purified dU-ssDNA of the 3Cs multiplex template plasmid, and the 3CS synthesis reactions were performed as described above.

### Generation of multiplexed 3Cs-gRNA libraries

#### 1. Equipment

Desktop microcentrifuge, shaking incubator at 37°C, 1.5 ml collection tubes, filtered sterile pipette tips, thermoblocks at 90°C and 50°C (e.g., Thermo Fisher, 88870004), an ultracentrifuge capable of spinning 50 ml falcon tubes at 10,000 rpm (Beckman Coulter Avanti J-30 I ultracentrifuge and a Beckman JA-12 fixed angle rotor), falcon tubes (polypropylene, 50 ml (Corning 352070)), a Bio-Rad Gene Pulser electroporation system (BioRad 164–2076), electroporation cuvettes Plus (2 mm, Model no. 620 (BTX)), a gel electrophoresis chamber, erlenmeyer flasks (glass, 200 ml and 500 ml), 10 cm dishes 10 cm plastic culture dishes (Corning, CLS430591), 14 ml round-bottom polystyrene tubes (e.g. Thermo Fisher, 10568531).

#### 2. dU-ssDNA template amplification

##### KCM transformation

KCM competent bacteria (Escherichia coli strain K12 CJ236, NEB, E4141) were transformed with 3Cs multiplex template plasmid template by mixing 100 ng of DNA with 2 µl of 5x KCM buffer (0.5M KCl, 0.15M CaCl2, 0.25M MgCl2) and water in a 10 µl reaction. After 10 min of incubation on ice, an equal volume of CJ236 bacteria was added to the DNA/KCM mixture, gently mixed, and chilled on ice for 15 min. The bacteria–DNA mixture was then incubated at room temperature for 10 min and subsequently inoculated into 200 µl of prewarmed SOC media (ThermoFisher Scientific, 15544034). Bacteria were incubated at 37°C and 200 rpm for 1 hr and then streaked on LB-agar plates with ampicillin (100 µg/ml) and chloramphenicol (34 µg/ml) for incubation overnight at 37°C.

##### Phage amplification and ssDNA purification

The morning after transformation, a single colony of transformed E. coli CJ236 was picked into 1 ml of 2YT media (Roth, 6676.2) supplemented with M13KO7 helper phage (NEB, N0315) to a final concentration of 1 × 10^8^ pfu/ml and ampicillin (final concentration 100 µg/ml) to maintain the host F′ episome and the phagemid, respectively. After 2 hrs of shaking at 200 rpm and 37°C, kanamycin (Roth, T832.3) was added to a final concentration of 25 µg/ml to select for bacteria that have been infected with M13KO7 helper phage. Bacteria were kept at 200 rpm and 37°C for 6 to 8 hrs. Afterwards, the culture was transferred to 30 ml of 2YT media supplemented with ampicillin (final concentration 100 µg/ml) and kanamycin (final concentration 25 µg/ml). After an additional 20 hrs of shaking at 200 rpm and 37°C, the bacterial culture was centrifuged for 10 min at 10,000 rpm and 4°C in a Beckman JA-12 fixed angle rotor. The supernatant was subsequently transferred to 6 ml (1/5 of culture volume) PEG/NaCl (20% polyethylene glycol 8,000, 2.5 M NaCl) and incubated for 1 hr at room temperature to precipitate phage particles. After 10 min of centrifugation at 10,000 rpm and 4°C in a Beckman JA-12 fixed angle rotor, the phage pellet was resuspended in 1.5 ml Dulbecco’s phosphate-buffered saline (PBS, Sigma, D8662) and centrifuged at 13,000 rpm for 5 min, before the phage-containing supernatant was transferred to a clean 1.5 ml microcentrifuge tube and stored at 4°C. Circular ssDNA was purified from the resuspended phages with the E.Z.N.A. M13 DNA Mini Kit (Omega Bio-Tek, D69001-01) according to the manufacturer’s protocol. Purity of the isolated ssDNA was ensured by agarose gel electrophoresis and purified ssDNA was stored at 4°C.

#### 3. Multiplexed 3Cs-DNA synthesis

The protocol for multiplexed 3Cs-DNA synthesis was adapted from Wegner et al., 2019 and optimized for reactions on the 3Cs multiplex template plasmid with two specific annealing sites. The oligonucleotides that were used for 3Cs reactions and the suppliers are listed separately (see ‘3Cs oligonucleotide design rules’ and ‘DNA oligonucleotides’).

##### Oligonucleotide phosphorylation and annealing

600 ng of oligonucleotides per annealing site (both, 3Cs h7SK- and hU6-oligonucleotides) were phosphorylated in two separate 20 µl reactions by mixing them with 2 µl 10x TM buffer (0.1 M MgCl2, 0.5 M Tris-HCl, pH 7.5), 2 µl 10 mM ATP (NEB, 0756), 1 µl 100 mM DTT (Cell Signaling Technology Europe, 7016), 20 units of T4 polynucleotide kinase (NEB, M0201) and water to a total volume of 20 µl. The mixture was incubated for 1 hr at 37°C. Phosphorylated oligonucleotides were immediately annealed to purified multiplex dU-ssDNA template by adding both 20 µl phosphorylation products to 25 µl 10x TM buffer, 20 µg of dU-ssDNA template and water to a total volume of 250 µl. The mixture was denatured for 5 min at 95°C, annealed for 5 min at 55°C and cooled down for 10 min at room temperature.

##### Multiplexed 3Cs-DNA reaction

3Cs-DNA was generated by adding 10 µl of 10 mM ATP, 10 µl of 100 mM dNTP mix (Roth, 0178.1/2), 15 µl of 100 mM DTT, 2000 ligation units of T4 DNA ligase (NEB, M0202), and 30 units of T7 DNA polymerase (NEB, M0274) to the annealed oligonucleotide–ssDNA mixture. The 3Cs synthesis mix was incubated for 12 hrs (overnight) at room temperature. Afterwards the 3Cs synthesis product was purified and desalted using a GeneJET Gel Extraction Kit (Thermo Fisher, K0692) according to the following protocol: 600 µl of binding buffer and 5 µl 3M sodium acetate (Sigma-Aldrich, 71196) were added to the synthesis product, mixed and applied to two purification columns, which were centrifuged for 3 min at 460 g. The flow-through was applied a second time to the same purification column to maximize yield. After two wash steps and 3 min of centrifugation at maximum speed, the DNA was eluted in 50 µl prewarmed water. The 3Cs reaction product was analyzed by gel electrophoresis alongside the dU-ssDNA template on a 0.8% TAE/agarose gel (100 V, 30 min).

#### 4. Multiplexed 3Cs-DNA library amplification, clean-up and quality control

##### Electroporation of 3Cs synthesis product

To amplify the multiplex 3Cs libraries, the 3Cs-DNA synthesis product was electroporated. To do so, 400 µl of electrocompetent E. coli (10-beta, NEB, C3020K) were thawed on ice and mixed with 6 µg of purified 3Cs-DNA. For electroporation, the DNA must be eluted in water or a low salt solution. After that, the mixture was incubated on ice for 15 min and then transferred into a cold 2 mm cuvette (BTX, 45–0125) that was then inserted into a Bio-Rad Gene Pulser with the following settings: resistance 200 Ω, capacity 25 F, voltage 2.5 kV. After electroporation, cells were rescued in 25 ml of pre-warmed SOC media and incubated for 30 min at 37°C and 200 rpm. After 30 min the culture was transferred into 400 ml of LB media supplemented with 100 μg/ml ampicillin.

##### Determination of transformation efficiency

To ensure library representation during and after amplification, the number of transformants was determined. After 30 min of incubation at 37°C and 200 rpm subsequently to the electroporation, a series of 10-fold dilutions of the 10 µl of bacterial culture in sterile Dulbecco’s phosphate-buffered saline (PBS, Sigma, D8662) was prepared. Dilutions were plated in triplicates on LB-agar containing 100 μg/ml ampicillin and incubated overnight at 37 °C. The next morning, the obtained colonies were counted. The number of transformants must be at least 100-fold higher than the library complexity to ensure maintenance of library diversity.

##### I-CeuI and I-SceI clean-up and quality control

Plasmid DNA of overnight liquid cultures was purified using the Qiagen Plasmid Maxi Kit (Qiagen, 12163), according to the manufacturer’s protocol to obtain the pre-library (P1). For removal of residual 3Cs template plasmid from the multiplex pre-library, 3 µg of purified DNA was digested with 10 units I-SceI (NEB, R0694), 10 units I-CeuI (NEB, R0699) and 5 µl NEB CutSmart buffer (NEB, B7204) in a reaction volume of 50 µl for 3 hrs at 37°C. After 3 hrs, an additional 10 units of I-SceI and I-CeuI and 5 µl NEB CutSmart buffer were added, as well as water to a final volume of 100 µl. After further incubation for additional 3 hours, the digestion reaction was subjected to gel electrophoresis on a 0.8% TAE/agarose gel (125 V, 40 min) to separate undigested 3Cs synthesis product from linearized template plasmid. The band resembling the undigested correct 3Cs synthesis product was purified using a Thermo Fisher Scientific GeneJET Gel Extraction Kit, according to the manufacturer’s protocol. Then, the purified 3Cs synthesis product was electroporated, according to the electroporation protocol described above. The next day, the resulting final 3Cs multiplex library preparation (P2) was purified from liquid culture using a Qiagen Plasmid Maxi Kit, according to the manufacturer’s protocol and quality controlled by analytical restriction enzyme digests, SANGER sequencing and by Next Generation Sequencing.

### Next-generation sequencing (NGS)

#### NGS sample preparation of 3Cs multiplex plasmid libraries

3Cs multiplex plasmid libraries were prepared for NGS as follows: 250 ng of plasmid DNA was used per PCR reaction and used in a volume of 50 µl using Next High-Fidelity 2x PCR Master Mix (NEB, M0541) (according to the manufacturer’s protocol), containing 2.5 µl of 10 µM primers each of forward and reverse primers. Depending on the library complexity up to four 50 µl reactions were performed. Primer sequences are listed separately (see ‘DNA oligonucleotides’). Thermal cycler parameters were set as follows: initial denaturation at 98°C for 5 min, 15 cycles of denaturation at 98°C for 30 s, annealing at 65°C for 30 s, extension at 72°C for 40 s, and final extension at 72°C for 5 min. PCR products were purified from a 1.5 % TAE/agarose gel using a GeneJet Gel Extraction Kit (Thermo Fisher Scientific), according to manufacturer’s protocol. Purified PCR products were denatured and diluted according to Illumina the guide lines and set to a final concentration of 2.6 pM in a total volume of 2.2 ml and 15% PhiX control and loaded onto a MiSeq, NextSeq 500 or NovaSeq sequencer (Illumina), according to manufacturer’s protocol. Sequencing was performed with single- or paired-end reads, 75 or 150 cycles, plus 8 cycles of index reading.

#### NGS sample preparation of 3Cs multiplex screening samples

To prepare 3Cs multiplex screening samples, the required amount of genomic DNA for sufficient coverage was calculated first: for the autophagy single and multiplex FACS screening samples, the required genomic DNA was calculated as *number of FACS sorted cells × screening coverage x* 6.6 *pg*. For the autophagy multiplex proliferation control screen, the required genomic DNA was determined by calculating *library complexity × screening coverage x* 6.6 *pg*.

The NGS sample preparation of all samples from screening with the biased libraries was performed with *library complexity ×* 200 (*maximum screening coverage*) *x* 6.6 *pg DNA*. The calculated amount of genomic DNA was used in a first PCR (PCR1) reaction with 2 to 4 µg of genomic DNA in a 50 µl reaction using the Next High-Fidelity 2x PCR Master Mix (NEB, M0541) (according to the manufacturer’s protocol) and 2.5 µl of 10 µM PCR1 primers, each of forward and reverse. Thermal cycler parameters were set as follows: initial denaturation at 98°C for 5 min, 15 cycles of denaturation at 98°C for 55 s, annealing at 65°C for 55 s, extension at 72°C for 110 s, and final extension at 72°C for 7 min. After PCR 1, 25 µl of PCR 1 product was transferred to a second PCR reaction (PCR2) in a 100 µl reaction with 50 µl High-Fidelity 2x PCR Master Mix and 2.5 µl of 10 µM PCR 2 primers that contain Illumina adaptors. Primer sequences for PCR1 and PCR2 are listed separately (see ‘DNA oligonucleotides’). Thermal cycler parameters were set as follows: initial denaturation at 98°C for 5 min, 10 cycles of denaturation at 98°C for 30 s, annealing at 65°C for 30 s, extension at 72°C for 40 s, and final extension at 72°C for 5 min. PCR products were purified from a 1.5 % TAE/agarose gel and processed for NGS sequencing as described for plasmid libraries.

### NGS data quality control and read count table generation

Raw next generation sequencing data were processed and demultiplexed with bcl2fastq v2.19.1.403 (Illumina). Read counts of individual gRNAs and gRNA combinations were determined using cutadapt 2.8, Bowtie2 2.3.0, and custom Python 3 scripts^67,68^. In brief, reads were trimmed with cutadapt using 5’ adapter sequences, truncated to 20 nucleotides, and aligned to the respective gRNA library using Bowtie2 with no mismatches allowed. The uniformity of each library distribution was assessed by plotting the cumulative distribution of all sequencing reads as a Lorenz curve and determining the area under the curve. The library distribution skew (skew ratio) of each library was determined by plotting the density of read counts and dividing the top 10 quantiles by the bottom 10 quantile. Cohen’s D statistics were applied to assess the quality of the biased libraries by comparing the distributions of non-targeting sequences and sequences targeting core essential genes^52^. Pairwise sample correlations were determined with Pearson’s correlation of the normalized read counts and visualized with hierarchically clustered heat maps (Seaborn library 0.10.1)^69^.

### Enrichment analyses

All enrichment analyses using MAGeCK were performed with median normalization of read counts and gRNAs with zero counts in the control samples were removed. Down-sampling of the 1:1 dataset was performed by randomly choosing 1 to 16 gRNA combinations per gene combination without replacement followed by individual MAGeCK analyses. gRNA combinations with an FDR≤10% and log2FC≤-0.5 were counted as statistically significant hits.

### Genetic interaction models

Interactions of gene pairs were computed according to five different models: SUM, MIN, LOG, MULT were used according to their definition in^17^, the MAX model defines the expected phenotype of a double gene-knockout as the maximal phenotype of the individual single gene-knockouts. The expected phenotypes of all double gene-knockouts were computed based on the phenotype of the respective single gene knockouts which were defined as the median log2FC (as provided by the MAGeCK analysis output) of all combinations of NHTs and gRNAs targeting the respective gene. For each model, the deviation of observed paired-knockout phenotypes from their expectation were expressed as their difference: Δlog2FC = observed - expected. Assuming that genetic interactions were rare, density plots of the dLFCs for each model were used to identify the model with the highest number of neutral interactions, indicated by a single large peak around 0 on the x-axis. Using the MAX model, we kept only combinations with p≤0.05 and a Δlog2FC larger than the standard deviation of all Δlog2FCs.

### Autophagy gene interaction network

To generate a network visualization based on our derived gene-gene interactions in autophagy, we exported MAX model-dependent delta log2FC per gene-gene interaction and imported them into the open source software platform for visualizing complex networks, Cytoscape (3.8.0)^70^. The style of the derived network was manually curated with the layout being set to a circular one.

### Cell culture

Cell culture was performed as described previously^45^. In brief, HEK293T cells (ATCC, CRL-3216) were maintained in Dulbecco’s Modified Eagle’s Medium (DMEM, Thermo Fisher Scientific, 41965–039) and puromycin-sensitive hTERT–RPE1 cells (provided by Andrew Holland) in DMEM: Nutrient Mixture F-12 (DMEM/F12, Thermo Fisher Scientific, 11320–074), each supplemented with 10% fetal bovine serum (FBS, Thermo Fisher Scientific, 10270) and 1% penicillin-streptomycin (Sigma-Aldrich, P4333) at 37°C with 5% CO2. In addition, hTERT–RPE1 cells were supplemented with 0.01 mg/ml hygromycin B (Capricorn Scientific, HYG-H). No method to ensure the state of authentication has been applied. Mycoplasma contamination testing was performed immediately after the arrival of the cells and multiple times during the course of the experiments. The hTERT-RPE1 GFP-LC3-RFP reporter cell line was generated by transducing hTERT-RPE1(Cas9) cells with retroviral particles generated with the transfer plasmid pMRX-IP-GFP-LC3-RFP (Addgene: 84573). Single cell clones were isolated and reporter functionality was tested by Torin1 and Bafilomycin A1 treatments.

### Genomic DNA extraction

Genomic DNA of cells was purified by resuspending PBS washed pellets of 40-50 million cells in 12 ml of TEX buffer (10 mM Tris-HCl, ph 7.5, 1 mM EDTA, ph 7.9, 0.5% SDS). Then 300 µl of proteinase K (10 mg/m) and 300 µl of Ribonuclease A (90 U/mg, 20 mg/ml) were added to the resuspended cells. The tubes were incubated overnight at 37°C at constant shaking. After complete cell lysis, 4 ml of 5 M NaCl was added, the solution was mixed and incubated at 4 °C for 40 min. After that the tubes were centrifuged at 14,000xg for 1 hr. The supernatant was transferred to a fresh tube and 24 ml of ice-cold 96% ethanol was added before the mixture was placed at -20°C overnight. The next day, the tubes were centrifuged at 14,000xg for 1 hr. Afterwards the supernatant was removed and the precipitated DNA was washed with ice-cold 70% ethanol. After further centrifugation 14,000xg for 1 hr the supernatant was removed and the DNA pellet was dried at room temperature and then dissolved in 5 ml of sterile water.

### GFP and mCherry knockouts

To examine the expression of gRNAs from both expression cassettes of the 3Cs multiplex template plasmid, two oligonucleotide pools were designed following the 3Cs oligonucleotide design rules. One oligonucleotide pool was designed for the h7SK cassette with 50 gRNAs targeting eGFP, the second pool was designed for the hU6 expression cassette with 50 gRNA targeting the mCherry gene. The two pools were used to generate three 3Cs libraries to selectively target either eGFP (GFP single library) or mCherry (mCherry single library) or both simultaneously (eGFP-mCherry multiplex library) following the protocol for generation of multiplexed 3Cs-gRNA libraries described above. Lentiviral supernatant of the three libraries was generated. Monoclonal hTERT–RPE1 cells with stable SpCas9, eGFP and mCherry expression were plated at 40% confluency. The next day, the cells were transduced with viral supernatant of one of the three libraries. After 48 hrs of transduction, cells were selected with puromycin (2.5µM) for 10 days. Then, eGFP and mCherry ratios were quantified by FACS analysis.

### Generation, quantification and transduction of lentiviral particles

Generation, quantification and transduction of lentiviral particles was performed as described previously^45^. In brief, the day before transfection, HEK293T cells were seeded to 2.5 × 10^5^ cells/ml. To transfect HEK293T cells, transfection media containing 1/10 of culture volume Opti-MEM I (Thermo Fisher Scientific, 31985–047), 10.5 µl Lipofectamin 2000 (Thermo Fisher Scientific, 11668019), 1.65 µg/ml transfer vector, 1.35 µg/ml pPAX2 (Addgene: 12260) and 0.5/ml µg pMD2.G (Addgene: 12259) was prepared. The mixture was incubated for 30 min at room temperature and added dropwise to the media. Lentiviral supernatant was harvested 48 hr after transfection and stored at −80°C.

To determine the lentiviral titer, hTERT–RPE1 cells were plated in a 6-well plate with 50,000 cells per well. The following day, cells were transduced in the presence of 8 µg/ml polybrene (Sigma, H9268) and a series of 0.5, 1, 5, and 10 µl of viral supernatant. After 2 days of incubation at 37°C, cells were subjected to puromycin selection for a total duration of 2 weeks, after which established colonies were counted per viral dilution. The number of colonies in the highest dilution was then volume normalized to obtain the final lentiviral titer.

To transduce hTERT–RPE1 cells, they were seeded at an appropriate density for each experiment with a maximal confluency of 60–70%. On the day of transduction, polybrene was added to the media to a final concentration of 8 µg/ml. The volume of lentiviral supernatant was calculated on the basis of the diversity of the respective library and of the desired coverage and multiplicity of infection (MOI) of the experiment. A MOI of 0.5 was applied to all screens. The number of cells that were transduced at the beginning of an experiment was calculated by multiplying the diversity of the library with the desired coverage and needed MOI.

### 3Cs CRISPR screening

#### Library distribution and experimental coverage interdependency screening

To explore the interdependency of multiplexed CRISPR library distribution and experimental screening coverage, three distorted 3Cs multiplex libraries were generated (see ‘generation of distorted libraries’) that represented libraries of different gRNA distributions. All three libraries were screened with a 20-fold and 200-fold coverage, each in triplicates. For the 20-fold screening, for each replicate, 1.1 million SpCas9 expressing hTERT–RPE1 cells were plated (0.37 million cells per flask) and transduced with the respective library with a MOI of 0.5. After 48 hrs the cells were selected with 2 µM puromycin and kept in growing conditions for 14 days. At day 14, the cells were harvested, pooled and stored at -20°C until their genomic DNA was extracted and processed for NGS. For the 200-fold screening a total of 11 million (0.5 million cells per flask) SpCas9 expressing hTERT–RPE1 cells were plated and transduced with the respective library with a MOI of 0.5. Further screening was performed identically to the 20-fold screen.

#### Combinatorial 3Cs-gRNA autophagy screening

Autophagy single and combinatorial gRNA screens for single or synergistic autophagy inhibition were performed in biological replicates and triplicates in the monoclonal hTERT–RPE1 cell line that stably expresses Streptococcus pyogenes Cas9 (SpCas9) and the autophagic flux probe (GFP-LC3-RFP)^56^, respectively. For each replicate 20 million cells (10 million for each, end time point and day 2 control) were transduced with lentiviral supernatant of the autophagy multiplex library with an MOI of 0.5 and a 1000- or 20-fold library coverage for single or combinatorial autophagy library screening, respectively. The control time points were harvested 2 days post-transduction. All remaining cells were kept in growing conditions until day 7, at what pint the cells were passaged, pooled and reseeded at library-diversity-maintaining density. After 13, 14 and 15 days the cells were treated with the mTOR inhibitor Torin1 (250 nM, InvivoGen, 1222998-36-8) for 24 hrs in three batches to induce autophagy. After 24 hrs of Torin1 treatment, cells were collected and 50,000 to 100,000 cells for single or 1.5 to 2.25 million cells for combinatorial screening of each batch were FACS sorted to enrich for cells with blocked autophagy. The sorted cells were reseeded and expanded for seven days before they were harvested, pooled and stored at -20°C until their genomic DNA was extracted and processed for NGS.

### FACS

Cell sorting was carried out with the FACS core facility of the Georg-Speyer Haus on a BD FACSAria Fusion, and CRISPR screening hit validation analysis on a FACSCanto II flow cytometer (BD Biosciences). Data was processed by FlowJo (FlowJo, LLC). Gating was carried out on the basis of viable and single cells that were identified on the basis of their scatter morphology.

### Arrayed autophagy candidate validation

The validation of single and combinatorial autophagy screening hits was performed in arrayed conditions (one knockout per well). To do so, single and dual gene-targeting CRISPR constructs were designed and generated. For each gene, the top scoring guide sequence was selected with Azimuth 2.0 of the GPP sgRNA Designer (https://portals.broadinstitute.org/gpp/public/analysis-tools/sgrna-design) and purchased as forward and reverse oligonucleotide with compatible overhangs for restriction enzyme cloning (see ‘DNA oligonucleotides’). The two oligonucleotides containing the gRNA target site were annealed and cloned into a restriction-enzyme digested and gel purified CRISPR vector. In more detail, single gRNA constructs were cloned into lenti-sgRNA blast vector (Addgene: 104993) by BsmBI restriction enzyme cloning. For combinatorial hit validation the dual CRISPR gRNA expression cassettes of pKLV2.2-h7SKgRNA5(SapI)-hU6gRNA5(BbsI)-PGKpuroBFP-W (Addgene: 72666) was cloned into the lenti-sgRNA blast plasmid to enable blasticidin selection of dual gRNA constructs. A silent point-mutation was introduced to remove the BbsI recognition site within the blasticidin sequence to allow the subsequent insertion of one gRNA by SapI (NEB, R0569) cloning into the h7SK expression cassette and the second gRNA by BbsI (NEB, R0539) cloning into the hU6 expression cassette. After cloning and sequence verification by SANGER sequencing, lentiviral supernatant was generated for each construct as described. Monoclonal hTERT–RPE1 cells with stable SpCas9 and GFP-LC3-RFP reporter expression were plated in 6-well plates with 50,000 cells per well. The following day, cells were transduced in the presence of 8 µg/ml polybrene (Sigma, H9268) with lentiviral supernatant. After 48 hrs the cells were selected with 10 µg/ml blasticidin (InvivoGen, ant-bl) for 7 days, passaged and cultivated at 40-60% confluency under constant blasticidin selection for an additional 7 days. At day 14, cells were treated with Torin1 to induce autophagy for 24 hrs until they were collected at day 15 and subject to FACS cell sorting to measure single or dual gene-knockout-induced autophagy blockage.

### gRNA performance and TIDE assay

Guide RNA performance was evaluated by TIDE assay, as described previously^45,71^. In short, for each gRNA sequence, +/- 400 nucleotides from the gRNA annealing site, PCR primers were designed to result in a PCR product of 800 to 1000 nts in length. The gRNA-locus is then PCR amplified with OneTaq DNA polymerase (NEB, M0480) using 1 µg of genomic DNA, 40 µM dNTPs (final concentration), 0.2 µM of each forward and reverse amplification primer, 10x OneTaq standard buffer, and 2.5 units of OneTaq DNA polymerase. PCR cycles were set up as follows: initial denaturation at 94°C for 3 min, 39 cycles of denaturation at 94°C for 20 s, annealing at 55°C for 30 s, strand extension at 68°C for 2 min, and final strand extension at 68°C for 5 min. The PCR products were analyzed on a 0.8% TAE/agarose gel (100 V, 30 min) and purified using a Thermo Fisher Scientific GeneJET Gel Extraction Kit according to the manufacturer’s protocol. The purified PCR product was pre-mixed with forward amplification primer and processed by SANGER sequencing, after which wildtype and gRNA-treated SANGER chromatograms were analyzed by TIDE and the percentage of unedited DNA extracted (https://tide.deskgen.com/).

### Data availability

NGS data are provided as raw read count tables as Supplementary Table 4. Plasmids encoding for extended- and combinatorial-autophagy libraries will be available through the Goethe University Depository (http://www.innovectis.de/INNOVECTIS-Frankfurt/Technologieangebote/Depository).

### Code availability

Custom software is publicly available from GitHuB, https://github.com/GEG-IBC2/3Cs-MPX (GEG-IBC2, 2019; copy archived at https://github.com/elifesciences-publications/3Cs).

### DNA oligonucleotides and oligo pools

Sequences of used DNA oligonucleotides and for 3Cs libraries are provided in Supplementary Table 2.

## FUNDING

Hessisches Ministerium für Wissenschaft und Kunst (IIIL5-518/17.004)

Hessisches Ministerium für Wissenschaft und Kunst (IIIL5-519/03/03.001)

Deutsche Forschungsgemeinschaft (EXC115/2)

Deutsche Forschungsgemeinschaft (EXC147/2)

Deutsche Forschungsgemeinschaft (EXC 2026)

Funders had no role in study design, data collection and interpretation, or the decision to submit the work for publication.

## ACKNOWLEDGEMENTS

We thank Andreas Ernst and Svenja Wiechmann for technical advice, Andrea Ballabio and Davide Cacchiarelli of the Telethon Institute of Genetics and Medicine, as well as Sebastian Wagner and Khalil Abou Elardat of the Cancer Genomics Core Facility Frankfurt, plus Tobias Schmidt of the Institute of Biochemistry I (Pathobiochemistry) for valuable support with NGS. This work was supported by the Hessian Ministry for Science and the Arts (HMWK, LOEWE-CGT, IIIL5-518/17.004), the German Research Foundation (DFG; CEF-MC - EXC115/2; ECCPS - EXC147/2; CPI - EXC 2026) and in part by the LOEWE Center Frankfurt Cancer Institute (FCI) funded by the Hessen State Ministry for Higher Education, Research and the Arts (IIIL5-519/03/03.001-0015).

## FIGURE LEGENDS

**Supp. Figure 1.**
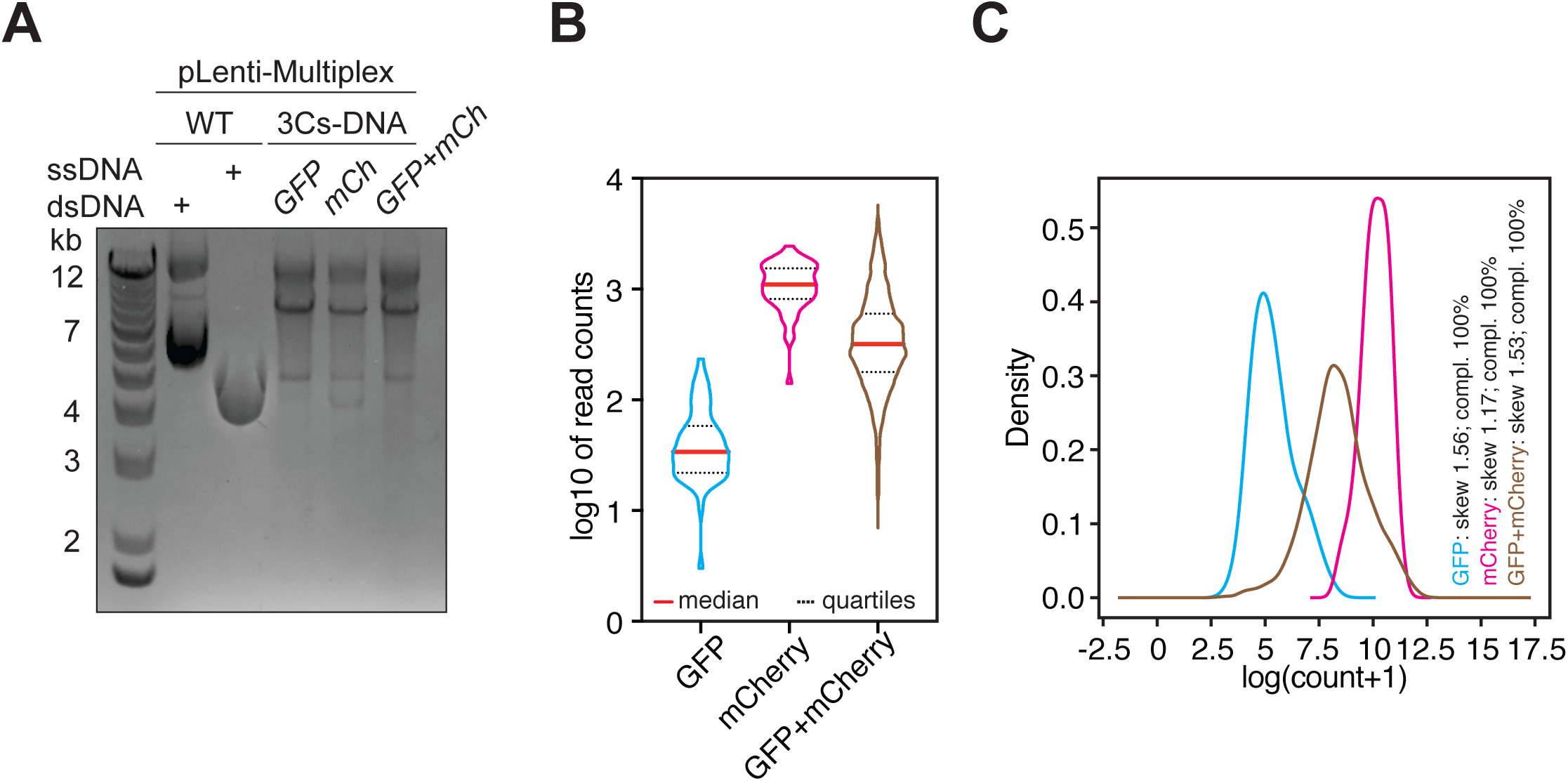
**A**) Analysis of single and multiplex dU-containing hetero-duplex 3Cs DNA by gel-electrophoresis. **B**) NGS sequencing depth of single and combinatorial GFP, mCherry, and GFP+mCherry 3Cs libraries. A sample’s median and quartiles are shown as red straight and black dotted line, respectively. **C**) Analysis of distribution skew (skew) and completeness (compl.) per library, based on read counts derived from B).

**Supp. Figure 2.**
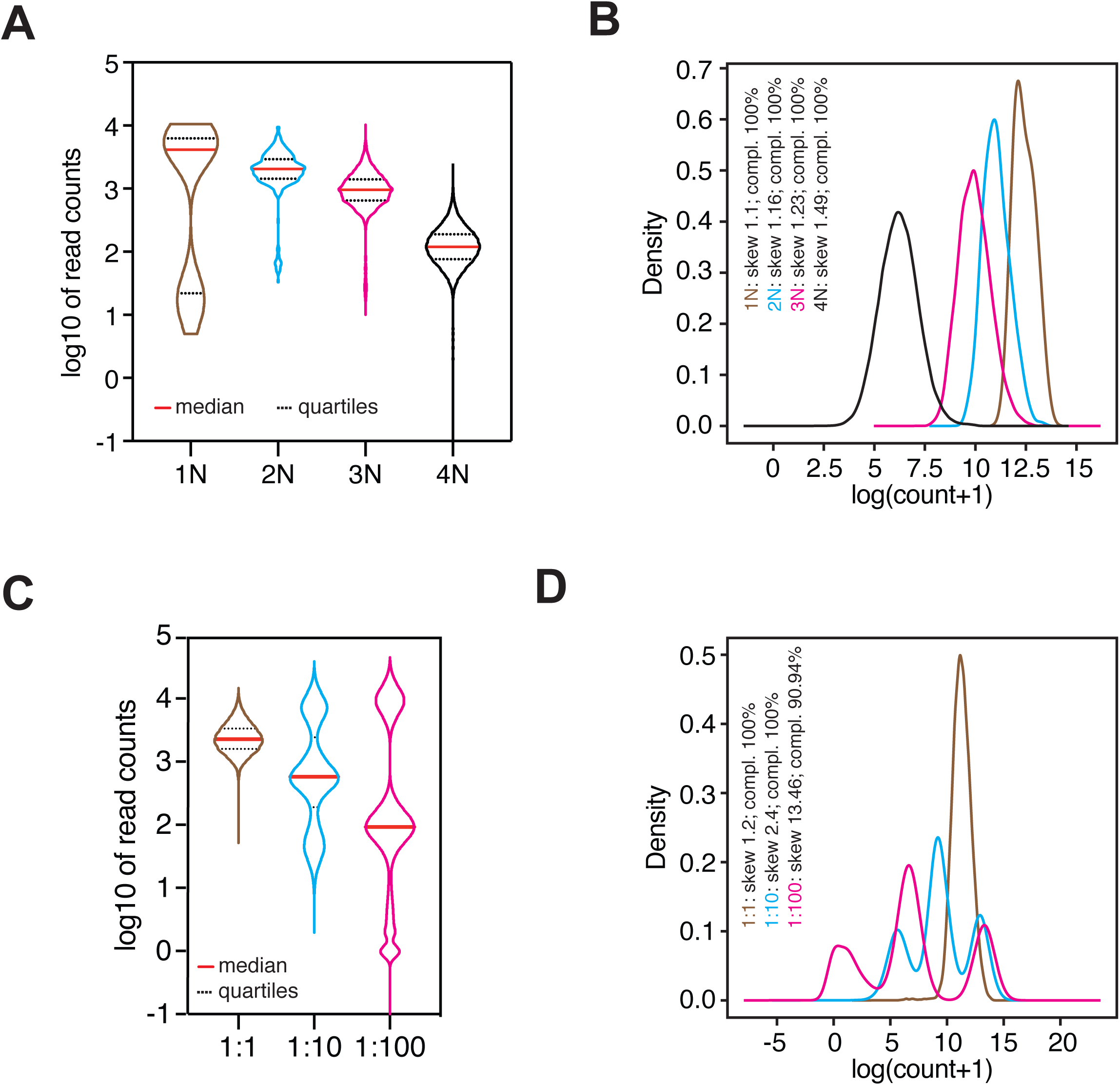
**A**) NGS sequencing depth of nucleotide-randomized combinatorial 3Cs libraries (1-4N). A library’s median and quartiles are shown as red straight and black dotted line, respectively. **B**) Analysis of distribution skew (skew) and completeness (compl.) per library, based on read counts derived from A). **C**) NGS sequencing depth of distribution skew biased combinatorial 3Cs libraries (1:1, 1:10, 1:100). A sample’s median and quartiles are shown as red straight and black dotted line, respectively. **D**) Analysis of distribution skew (skew) and completeness (compl.) per library, based on read counts derived from C).

**Supp. Figure 3.**
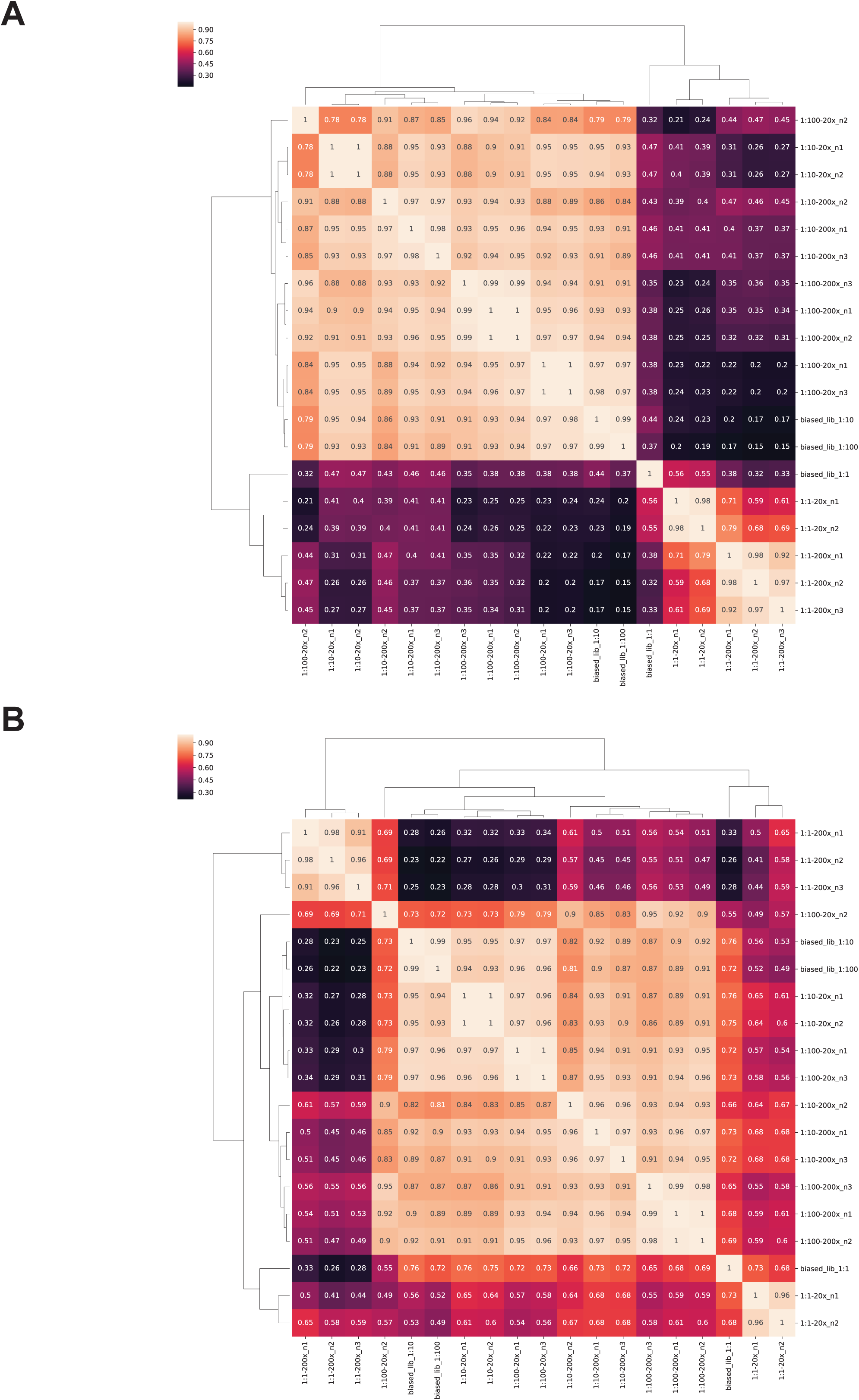
**A-B**) Pairwise sample correlation (Pearson’s correlation coefficient), visualized as hierarchically clustered heatmaps (n), library distributions skews (1:1, 1:10, 1:100) and coverages (20x, 200x) on gRNA-pair (A) and gene-pair (B) levels. Color code based on Pearson’s correlation coefficient (rho) of normalized gRNA read counts.

**Supp. Figure 4.**
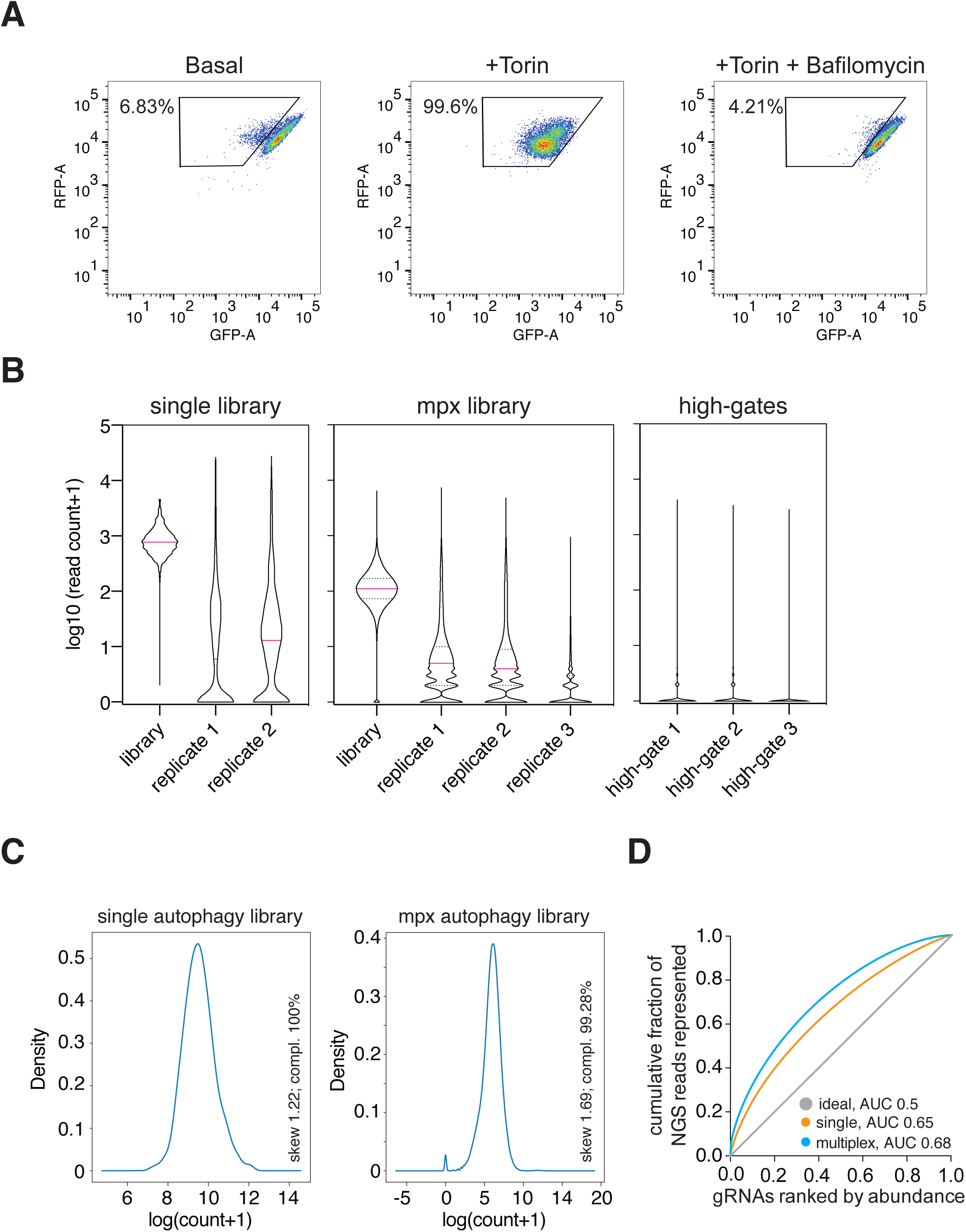
**A**) FACS analysis of monoclonal hTERT-RPE1 GFP-LC3B-RFP reporter cell line under conditions of basal autophagy (Basal), Torin1-induced autophagy (+Torin1), and Torin1-induced but Bafilomycin A1-blocked autophagy (+Torin1 + Bafilomycin A1). Gating is based on Torin1-induced reduction of GFP signal; percentage (%) of cells in gate. **B**) Analysis of NGS sequencing depth per 3Cs library (single, mpx) and replicate post-FACS sample (1-3). A sample’s median and quartiles are shown as red straight and black dotted line, respectively. **C**) Analysis of distribution skew (skew) and completeness (compl.) per autophagy library (single, mpx), based on read counts derived from B). **D**) Area-under-the-curve (AUC) determination of the single and combinatorial (multiplex) autophagy libraries. As a reference, a perfectly distributed library (ideal) is shown in grey. AUC values are indicated next to each library’s identifier.

**Supp. Figure 5.**
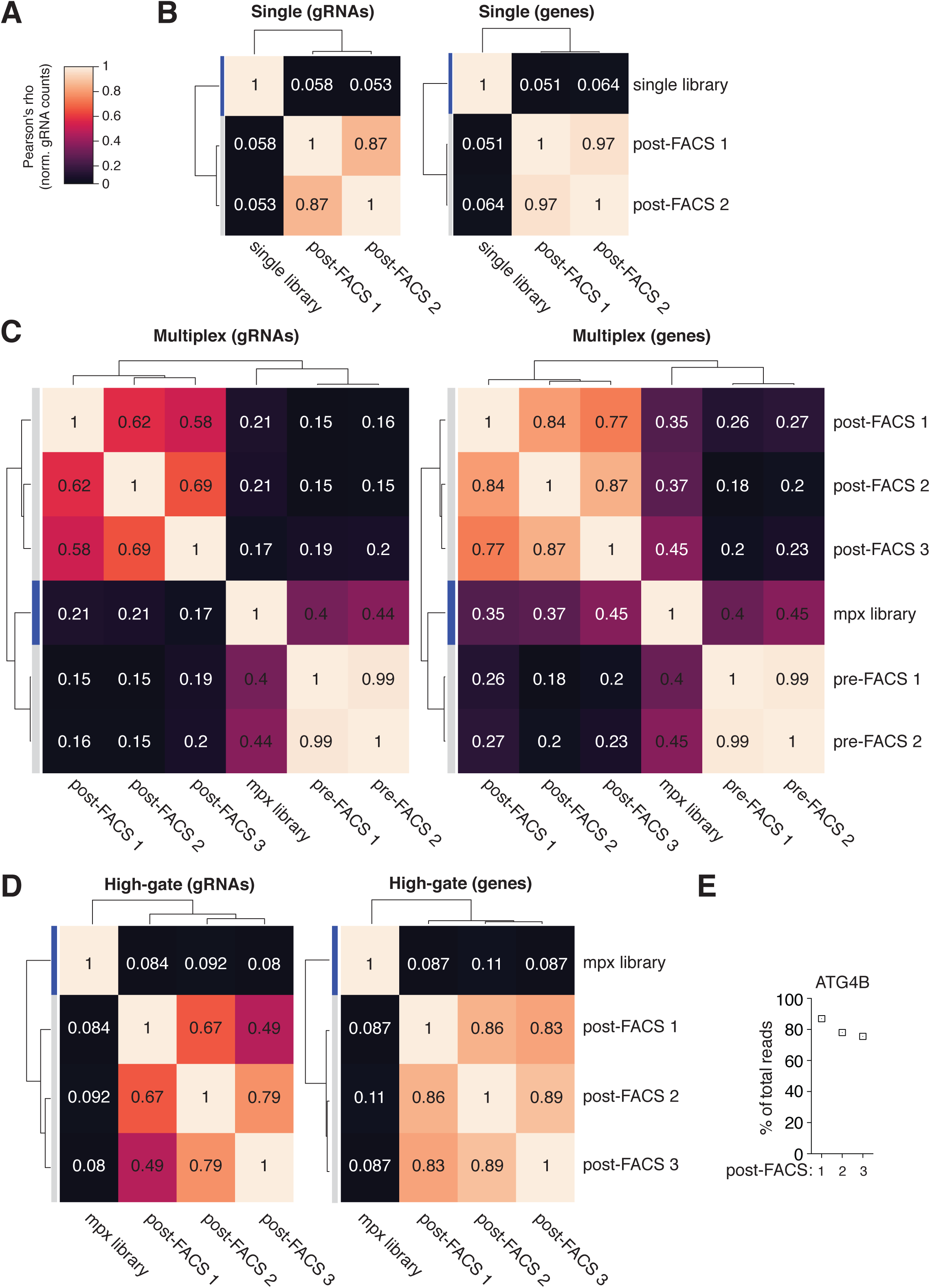
**A-D**) Pairwise sample correlation (Pearson’s correlation coefficient), visualized as hierarchically clustered heatmaps (1-3) of pre- and post-FACS samples of autophagy enrichment screens with single (B) and combinatorial gRNA-targeting in autophagy blockage (C) and high-gates (D) on gRNA and gene level. Color code based on Pearson’s correlation coefficient (rho) of normalized gRNA read counts. **E**) Analysis of ATG4B-associated guide pairs in high-gate post-FACS samples.

**Supp. Figure 6.**
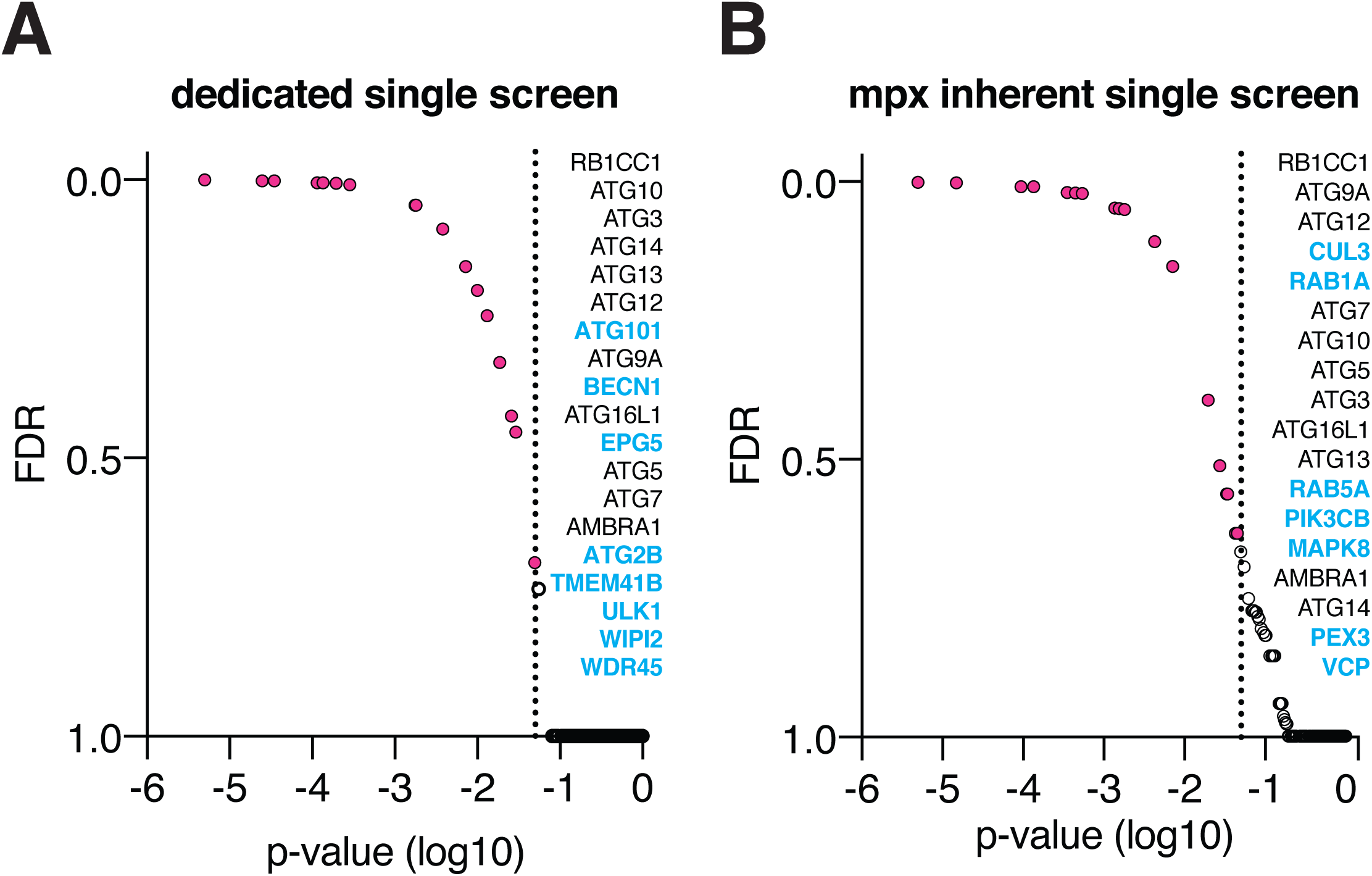
**A-B**) MAGeCK analysis of dedicated single (A) and multiplex-inherent single (B) gRNA autophagy screens between pre- and post-FACS samples with genes in red when matching cutoff criteria of p-value≤0.05. Screen-selective hits in blue.

**Supp. Figure 7.**
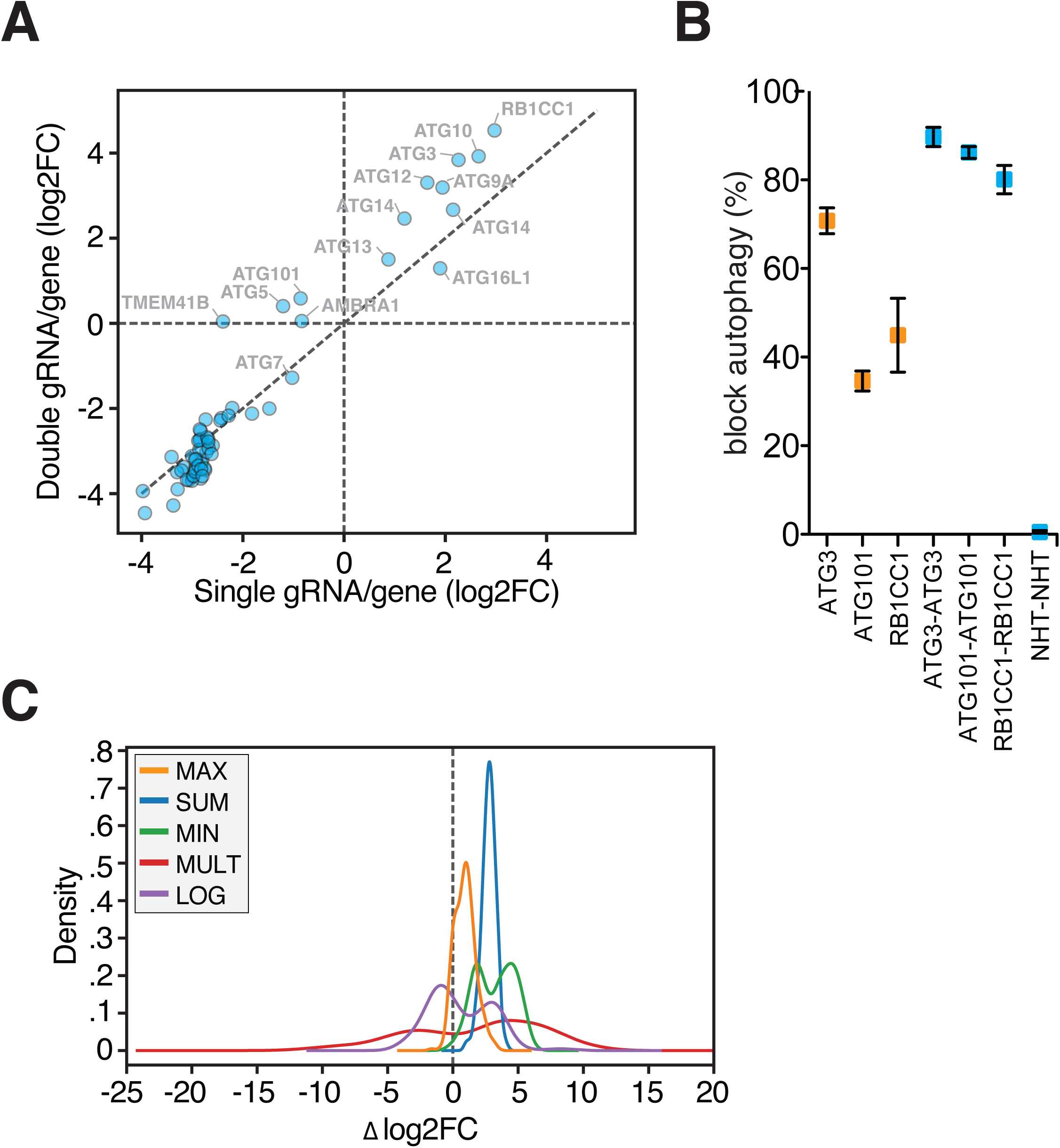
**A**) Log2FC-analysis of targeting a single gene with one (single) or two (double) gRNAs. The dotted diagonal line represents equal phenotypic strength, based on FACS enrichment. **B**) Arrayed analysis of phenotypic strength of single essential genes for autophagy when targeted with one (yellow) or two (blue) gRNAs. Error bars represent standard error of mean (SEM) over three biological replicates (n=3). **C**) Density plots of delta log2FC (Δlog2FC) value analyses of single essential genes for autophagy, computed by MAX, SUM, MIN, MULT, and LOG models.

## REFERENCES

1. Maddalo, D. et al. In vivo engineering of oncogenic chromosomal rearrangements with the CRISPR/Cas9 system. Nature 516, 423–428 (2014).

2. Cong, L. et al. Multiplex genome engineering using CRISPR/Cas systems. Science (80-.). 339, 819–823 (2013).

3. Wong, A. S. L. et al. Multiplexed barcoded CRISPR-Cas9 screening enabled by CombiGEM. Proc. Natl. Acad. Sci. U. S. A. 113, 2544–2549 (2016).

4. Tzelepis, K. et al. A CRISPR Dropout Screen Identifies Genetic Vulnerabilities and Therapeutic Targets in Acute Myeloid Leukemia. Cell Rep 17, 1193–1205 (2016).

5. Kabadi, A. M., Ousterout, D. G., Hilton, I. B. & Gersbach, C. A. Multiplex CRISPR/Cas9- based genome engineering from a single lentiviral vector. Nucleic Acids Res. 42, e147–e147 (2014).

6. Sakuma, T., Nishikawa, A., Kume, S., Chayama, K. & Yamamoto, T. Multiplex genome engineering in human cells using all-in-one CRISPR/Cas9 vector system. Sci Rep 4, 5400 (2014).

7. Vad-Nielsen, J., Lin, L., Bolund, L., Nielsen, A. L. & Luo, Y. Golden Gate Assembly of CRISPR gRNA expression array for simultaneously targeting multiple genes. Cell. Mol. Life Sci. 73, 4315–4325 (2016).

8. Zuckermann, M. et al. A novel cloning strategy for one-step assembly of multiplex CRISPR vectors. Sci. Rep. 8, (2018).

9. Albers, J. et al. A versatile modular vector system for rapid combinatorial mammalian genetics. J. Clin. Invest. 125, 1603–1619 (2015).

10. Haldeman, J. M. et al. Creation of versatile cloning platforms for transgene expression and dCas9-based epigenome editing. Nucleic Acids Res. 47, e23–e23 (2018).

11. Breunig, C. T. et al. One step generation of customizable gRNA vectors for multiplex CRISPR approaches through string assembly gRNA cloning (STAgR). PLoS One 13, (2018).

12. Nissim, L., Perli, S. D., Fridkin, A., Perez-Pinera, P. & Lu, T. K. Multiplexed and Programmable Regulation of Gene Networks with an Integrated RNA and CRISPR/Cas Toolkit in Human Cells. Mol. Cell 54, 698–710 (2014).

13. Minkenberg, B., Wheatley, M. & Yang, Y. Chapter Seven - CRISPR/Cas9-Enabled Multiplex Genome Editing and Its Application. in Gene Editing in Plants (eds. Weeks, D. P. & Yang, B. B. T.-P. in M. B. and T. S.) 149, 111–132 (Academic Press, 2017).

14. Vidigal, J. A. & Ventura, A. Rapid and efficient one-step generation of paired gRNA CRISPR-Cas9 libraries. Nat Commun 6, 8083 (2015).

15. Shen, J. P. et al. Combinatorial CRISPR-Cas9 screens for de novo mapping of genetic interactions. Nat. Methods 14, 573–576 (2017).

16. Han, K. et al. Synergistic drug combinations for cancer identified in a CRISPR screen for pairwise genetic interactions. Nat. Biotechnol. 35, 463–474 (2017).

17. Mani, R., St Onge, R. P., Hartman, J. L., Giaever, G. & Roth, F. P. Defining genetic interaction. Proc. Natl. Acad. Sci. U. S. A. 105, 3461–6 (2008).

18. Nijman, S. M. B. Synthetic lethality: General principles, utility and detection using genetic screens in human cells. FEBS Letters 585, 1–6 (2011).

19. Chan, D. A. & Giaccia, A. J. Harnessing synthetic lethal interactions in anticancer drug discovery. Nature Reviews Drug Discovery 10, 351–364 (2011).

20. Ramkumar, P., Kampmann, M. & Qian, C. CRISPR-based genetic interaction maps inform therapeutic strategies in cancer. Translational Cancer Research 7, S61–S67 (2018).

21. Ashworth, A. A Synthetic Lethal Therapeutic Approach: Poly(ADP) Ribose Polymerase Inhibitors for the Treatment of Cancers Deficient in DNA Double-Strand Break Repair. J. Clin. Oncol. 26, 3785–3790 (2008).

22. Helleday, T. The underlying mechanism for the PARP and BRCA synthetic lethality: Clearing up the misunderstandings. Molecular Oncology 5, 387–393 (2011).

23. Cho, S. Y. et al. A novel combination treatment targeting BCL-XL and MCL1 for KRAS/BRAF-mutated and BCL2L1-amplified colorectal cancers. Mol. Cancer Ther. 16, 2178–2190 (2017).

24. Najm, F. J. et al. Orthologous CRISPR–Cas9 enzymes for combinatorial genetic screens. Nat. Biotechnol. 36, 179–189 (2017).

25. Mereniuk, T. R. et al. Synthetic lethal targeting of PTEN-deficient cancer cells using selective disruption of polynucleotide kinase/phosphatase. Mol. Cancer Ther. 12, 2135–2144 (2013).

26. Neshat, M. S. et al. Enhanced sensitivity of PTEN-deficient tumors to inhibition of FRAP/mTOR. Proc. Natl. Acad. Sci. U. S. A. 98, 10314–10319 (2001).

27. Zamanighomi, M. et al. GEMINI: a variational Bayesian approach to identify genetic interactions from combinatorial CRISPR screens. Genome Biol. 20, 137 (2019).

28. Klionsky, D. J. & Emr, S. D. Autophagy as a regulated pathway of cellular degradation. Science 290, 1717–1721 (2000).

29. Dikic, I. & Elazar, Z. Mechanism and medical implications of mammalian autophagy. Nature Reviews Molecular Cell Biology 19, 349–364 (2018).

30. Mizushima, N., Yoshimori, T. & Levine, B. Methods in Mammalian Autophagy Research. Cell 140, 313–326 (2010).

31. Yoshii, S. R. & Mizushima, N. Monitoring and measuring autophagy. International Journal of Molecular Sciences 18, (2017).

32. Morita, K. et al. Genome-wide CRISPR screen identifies TMEM41B as a gene required for autophagosome formation. J Cell Biol 217, 3817–3828 (2018).

33. Shoemaker, C. J. et al. CRISPR screening using an expanded toolkit of autophagy reporters identifies TMEM41B as a novel autophagy factor. PLoS Biol. 17, e2007044 (2019).

34. Moretti, F. et al. TMEM 41B is a novel regulator of autophagy and lipid mobilization. EMBO Rep. 19, (2018).

35. Jia, R. & Bonifacino, J. S. Negative Regulation of Autophagy by UBA6-BIRC6–Mediated Ubiquitination of LC3. bioRxiv 8, 699124 (2019).

36. Orvedahl, A. et al. Autophagy genes in myeloid cells counteract IFNγ-induced TNF- mediated cell death and fatal TNF-induced shock. Proc. Natl. Acad. Sci. U. S. A. 116, 16497–16506 (2019).

37. Kerins, M. J. et al. Genome-Wide CRISPR Screen Reveals Autophagy Disruption as the Convergence Mechanism That Regulates the NRF2 Transcription Factor. Mol. Cell. Biol. 39, (2019).

38. Towers, C. G. et al. Cancer Cells Upregulate NRF2 Signaling to Adapt to Autophagy Inhibition. Dev. Cell 50, 690-703.e6 (2019).

39. Potting, C. et al. Genome-wide CRISPR screen for PARKIN regulators reveals transcriptional repression as a determinant of mitophagy. Proc. Natl. Acad. Sci. 115, 201711023 (2017).

40. Hoshino, A. et al. The ADP/ATP translocase drives mitophagy independent of nucleotide exchange. Nature 575, 375–379 (2019).

41. Heo, J. M. et al. Integrated proteogenetic analysis reveals the landscape of a mitochondrial-autophagosome synapse during PARK2-dependent mitophagy. Sci. Adv. 5, (2019).

42. Towers, C. G. & Thorburn, A. Therapeutic Targeting of Autophagy. EBioMedicine 14, 15–23 (2016).

43. Levy, J. M. M., Towers, C. G. & Thorburn, A. Targeting autophagy in cancer. Nature Reviews Cancer 17, 528–542 (2017).

44. Galluzzi, L., Bravo-San Pedro, J. M., Levine, B., Green, D. R. & Kroemer, G. Pharmacological modulation of autophagy: Therapeutic potential and persisting obstacles. Nature Reviews Drug Discovery 16, 487–511 (2017).

45. Wegner, M. et al. Circular synthesized CRISPR/Cas gRNAs for functional interrogations in the coding and noncoding genome. Elife 8, (2019).

46. Dang, Y. et al. Optimizing sgRNA structure to improve CRISPR-Cas9 knockout efficiency. Genome Biol. 16, 280 (2015).

47. Cross, B. C. S. et al. Increasing the performance of pooled CRISPR-Cas9 drop-out screening. Sci. Rep. 6, 1–8 (2016).

48. Huang, R., Fang, P. & Kay, B. K. Improvements to the Kunkel mutagenesis protocol for constructing primary and secondary phage-display libraries. Methods 58, 10–17 (2012).

49. Imkeller, K., Ambrosi, G., Boutros, M. & Huber, W. Gscreend: Modelling asymmetric count ratios in CRISPR screens to decrease experiment size and improve phenotype detection. Genome Biol. 21, 53 (2020).

50. Hart, T. et al. High-Resolution CRISPR Screens Reveal Fitness Genes and Genotype- Specific Cancer Liabilities. Cell 163, 1515–1526 (2015).

51. Doench, J. G. et al. Optimized sgRNA design to maximize activity and minimize off-target effects of CRISPR-Cas9. Nat Biotechnol 34, 184–191 (2016).

52. Kim, E. & Hart, T. Improved analysis of CRISPR fitness screens and reduced off-target effects with the BAGEL2 gene essentiality classifier. bioRxiv 2020.05.30.125526 (2020). doi:10.1101/2020.05.30.125526

53. Li, W. et al. MAGeCK enables robust identification of essential genes from genome-scale CRISPR/Cas9 knockout screens. Genome Biol. 15, 554 (2014).

54. Gonatopoulos-Pournatzis, T. et al. Genetic interaction mapping and exon-resolution functional genomics with a hybrid Cas9–Cas12a platform. Nat. Biotechnol. 38, 638–648 (2020).

55. Ong, S. H., Li, Y., Koike-Yusa, H. & Yusa, K. Optimised metrics for CRISPR-KO screens with second-generation gRNA libraries. Sci. Rep. 7, 7384 (2017).

56. Kaizuka, T. et al. An Autophagic Flux Probe that Releases an Internal Control. Mol. Cell 64, 835–849 (2016).

57. Dejesus, R. et al. Functional CRISPR screening identifies the ufmylation pathway as a regulator of SQSTM1/p62. Elife 5, (2016).

58. Goodwin, J. M. et al. Autophagy-Independent Lysosomal Targeting Regulated by ULK1/2-FIP200 and ATG9. Cell Rep. 20, 2341–2356 (2017).

59. Peets, E. M. et al. Minimized double guide RNA libraries enable scale-limited CRISPR/Cas9 screens. bioRxiv 859652 (2019). doi:10.1101/859652

60. Costanzo, M. et al. A global genetic interaction network maps a wiring diagram of cellular function. Science 353, aaf1420 (2016).

61. Xie, X., Le, L., Fan, Y., Lv, L. & Zhang, J. Autophagy is induced through the ROS-TP53- DRAM1 pathway in response to mitochondrial protein synthesis inhibition. Autophagy 8, 1071–1084 (2012).

62. Gonçalves, E. et al. Minimal genome-wide human CRISPR-Cas9 library. bioRxiv 848895 (2019). doi:10.1101/848895

63. Dede, M., McLaughlin, M., Kim, E. & Hart, T. Multiplex enCas12a screens show functional buffering by paralogs is systematically absent from genome-wide CRISPR/Cas9 knockout screens. bioRxiv 2020.05.18.102764 (2020). doi:10.1101/2020.05.18.102764

64. Folkerts, H. et al. Inhibition of autophagy as a treatment strategy for p53 wild-type acute myeloid leukemia. Cell Death Dis. 8, e2927 (2017).

65. Rogov, V., Dötsch, V., Johansen, T. & Kirkin, V. Interactions between Autophagy Receptors and Ubiquitin-like Proteins Form the Molecular Basis for Selective Autophagy. Molecular Cell 53, 167–178 (2014).

66. Chen, B. et al. Dynamic imaging of genomic loci in living human cells by an optimized CRISPR/Cas system. Cell 155, 1479–1491 (2013).

67. Martin, M. Cutadapt removes adapter sequences from high-throughput sequencing reads. EMBnet.journal 17, 10 (2011).

68. Langmead, B. & Salzberg, S. L. Fast gapped-read alignment with Bowtie 2. Nat. Methods 9, 357–359 (2012).

69. Waskom, M. et al. mwaskom/seaborn: v0.10.1 (April 2020). (2020). doi:10.5281/ZENODO.3767070

70. Shannon, P. et al. Cytoscape: A software Environment for integrated models of biomolecular interaction networks. Genome Res. 13, 2498–2504 (2003).

71. Brinkman, E. K., Chen, T., Amendola, M. & van Steensel, B. Easy quantitative assessment of genome editing by sequence trace decomposition. Nucleic Acids Res 42, e168 (2014).

